# A serine-folate metabolic unit controls resistance and tolerance of infection

**DOI:** 10.1101/2022.11.25.517956

**Authors:** Krista Grimes, Esteban J. Beckwith, William H. Pearson, Jake Jacobson, Surabhi Chaudhari, Gabriel N. Aughey, Gerald Larrouy-Maumus, Tony D. Southall, Marc S. Dionne

**Author notes:** These authors contributed equally (Krista Grimes, Esteban J. Beckwith).

## Abstract

Immune activation drives metabolic change in most animals. Immune-induced metabolic change is most conspicuous as a driver of pathology in serious or prolonged infection, but it is normally expected to be important to support immune function and recovery. Many of the signalling mechanisms linking immune detection with metabolic regulation, and their specific consequences, are unknown. Here, we show that *Drosophila melanogaster* respond to many bacterial infections by altering expression of genes of the folate cycle and associated enzymes of amino acid metabolism. The net result of these changes is increased flow of carbon from glycolysis into serine and glycine synthesis and a shift of folate cycle activity from the cytosol into the mitochondrion. Immune-induced transcriptional induction of *astray* and *Nmdmc*, the two most-induced of these enzymes, depends on *Dif* and *foxo*. Loss of *astray* or *Nmdmc* results in infection-specific immune defects. Our work thus shows a key mechanism that connects immune-induced changes in metabolic signalling with the serine-folate metabolic unit to result in changed immune function.

## Introduction

Immunity and metabolism are closely interconnected. Immune responses are metabolically costly, and therefore metabolism must change to provide the required energy and raw materials. As a result of this dependence, malnutrition and other metabolic perturbations can negatively impact immune function^1^. Conversely, inappropriate or prolonged immune activation can drive metabolic pathology, exemplified in the dramatic wasting of lean and fatty tissues in tuberculosis patients^2^. Malnourished children have defects in both innate and adaptive immunity, increasing their risk of dying from infection^3^, whilst obesity is associated with chronic inflammation which in turn promotes the development of other metabolic disorders^4^. Despite the importance of immune-metabolic interplay in human disease, we do not yet fully understand the mechanisms and consequences of this interaction.

As a genetically tractable organism with similar innate immunity and metabolism to humans, *Drosophila melanogaster* has been widely used to study immune-metabolic interactions *in vivo*^5,6^. The fat body, an organ analogous to the liver and adipose tissue of vertebrates, is the primary site of energy storage and metabolism in the fly^7^. However, it is also responsible for coordinating the humoral immune response. Activation of Toll or Imd signalling in the fat body, triggered most commonly by Gram-positive or -negative bacteria respectively, results in cytosolic translocation of the NF-κB transcription factors Dif and Relish^8^. These drive expression antimicrobial peptides (AMPs), which are secreted from the fat body and neutralize microorganisms via a variety of mechanisms^8^.

Infection can profoundly affect fat body energy storage. A cachexia-like phenotype has been observed upon *Mycobacterium marinum* infection, in which flies progressively lose glycogen and triglyceride stores and become hyperglycemic^9^. Similar observations have been made during *Listeria monocytogenes*^10^, *Micrococcus luteus*^11^ and *Beauvaria bassiana*^11^ infection. This reduction in energy stores is likely at least in part a direct result of immune signalling, as constitutive activation of the Toll receptor in the larval fat body phenocopies triglyceride loss^11^.

One mechanism by which the Toll pathway can drive changes in triglyceride and carbohydrate metabolism is through disruption of insulin signalling, a prominent anabolism-promoting pathway. Insulin signalling results in the phosphorylation and activation of Akt, which inhibits the transcription factor FOXO^12^. Infections with *M. marinum, M. luteus and B. bassiana* all reduce phosphorylation of Akt^9,11^. Overexpression of Dif also reduces Akt phosphorylation^11^, indicating that disruption of the insulin signalling pathway likely occurs via transcriptional targets of Dif. The mechanism of disruption has been attributed to impaired Akt phosphorylation by the kinase Pdk1^13^, and reduced insulin like peptide activity^14^. This reduction in insulin pathway activity has been suggested to enable triglyceride remodelling into phospholipids to support the immune response^15^. Outside of insulin signalling, the transcription factor MEF2 has also been shown to be an important determinant of triglyceride and carbohydrate anabolic-catabolic balance in infection^16^.

Whilst a significant amount is known in *Drosophila* about the mechanisms connecting immune activation with triglyceride and carbohydrate metabolism, connections between immune activation and other metabolic processes are not as clearly understood. One such area is amino acid metabolism. Two studies have noted that immune activation drives changes to the expression of enzymes involved in amino acid metabolism. Infection of *Drosophila* with seven different bacteria results in downregulation of *Agxt*, encoding an alanine-glyoxylate aminotransferase involved in alanine and glycine metabolism^17^. Further, constitutive activation of Imd in the larval fat body has been shown to drive upregulation of genes encoding enzymes associated with glycine, serine and threonine metabolism, and downregulation of arginine and proline metabolism^18^.

Beyond changes to transcript levels, two studies have measured the levels of amino acids themselves and noted infection-induced changes. Both *Candida albicans* and *L. monocytogenes* infections result in a decrease in branched chain amino acids^10,19^. *L. monocytogenes* infection causes changes to the levels of numerous other amino acids, including a consistent increase in amino acids associated with glycine, serine and threonine metabolism, and a drop in tyrosine^10^. The reduction in tyrosine is likely due to its involvement in generating phenols for melanisation, an important aspect of the immune response^10,20^. However, the functional impacts of alterations to the other individual amino acids, and the expression of associated enzymes, remains unexplored.

This is also true of other metabolic systems that support amino acid metabolism. For example, one-carbon (1C) metabolism encompasses a set of reactions that involve the activation and transfer of one-carbon units via folate intermediates^21^. Carbon units are moved between metabolites to support biosynthetic processes such as purine and pyrimidine biosynthesis, as well as maintain glycine, serine, and methionine homeostasis^21^. Infection-induced alterations to the expression of enzymes involved in 1C metabolism have been observed in *Drosophila*^17,18^, but the underlying regulatory mechanisms and functional relevance are unknown.

In this work, we performed RNAseq and fat body-specific Targeted DamID (TaDa) to study gene expression changes induced by infection. We observed transcriptional upregulation of two enzymes of interest—*astray*, a phosphoserine phosphatase, and *Nmdmc*, an enzyme involved in 1C metabolism that supplies a tetrahydrofolate cofactor for serine to glycine conversion. We observed that other enzymes in metabolic proximity are also transcriptionally regulated during infection, including *Shmt*, which catalyses the serine to glycine conversion, and *pug*, another 1C enzyme that is partially redundant with *Nmdmc*. We first looked at regulation of the enzymes and discovered that a transcriptional preference is shown towards running the 1C cycle through the mitochondria upon infection. We found that *astray* and *Nmdmc* are repressed by MEF2 in healthy flies and upregulated by Dif and FOXO, perhaps via the suppression of insulin signalling. Finally, we used knock downs and CRISPR mutants to investigate the functional role of these enzymes during infection. We found that *astray* is required for tolerance during *Staphylococcus aureus* infection and resistance against *Enterococcus faecalis*, whilst the other enzymes show mixed, pathogen specific phenotypes.

## Materials & Methods

### Fly husbandry

Flies were maintained on a standard yeast-sugar diet containing 10% w/v Brewer’s yeast, 8% fructose, 2% polenta, 0.8% agar, 0.075% nipagin and 0.0825% v/v propanoic acid. Flies were kept in a 12-hour light-dark cycle at 25°C unless indicated otherwise. The genetic lines used in this study are presented in Table 1. Any lines used in experiments were moved onto our *w^1118^* genetic background using isogenic balancer chromosomes. All experimental flies were male.

**Table 1.**
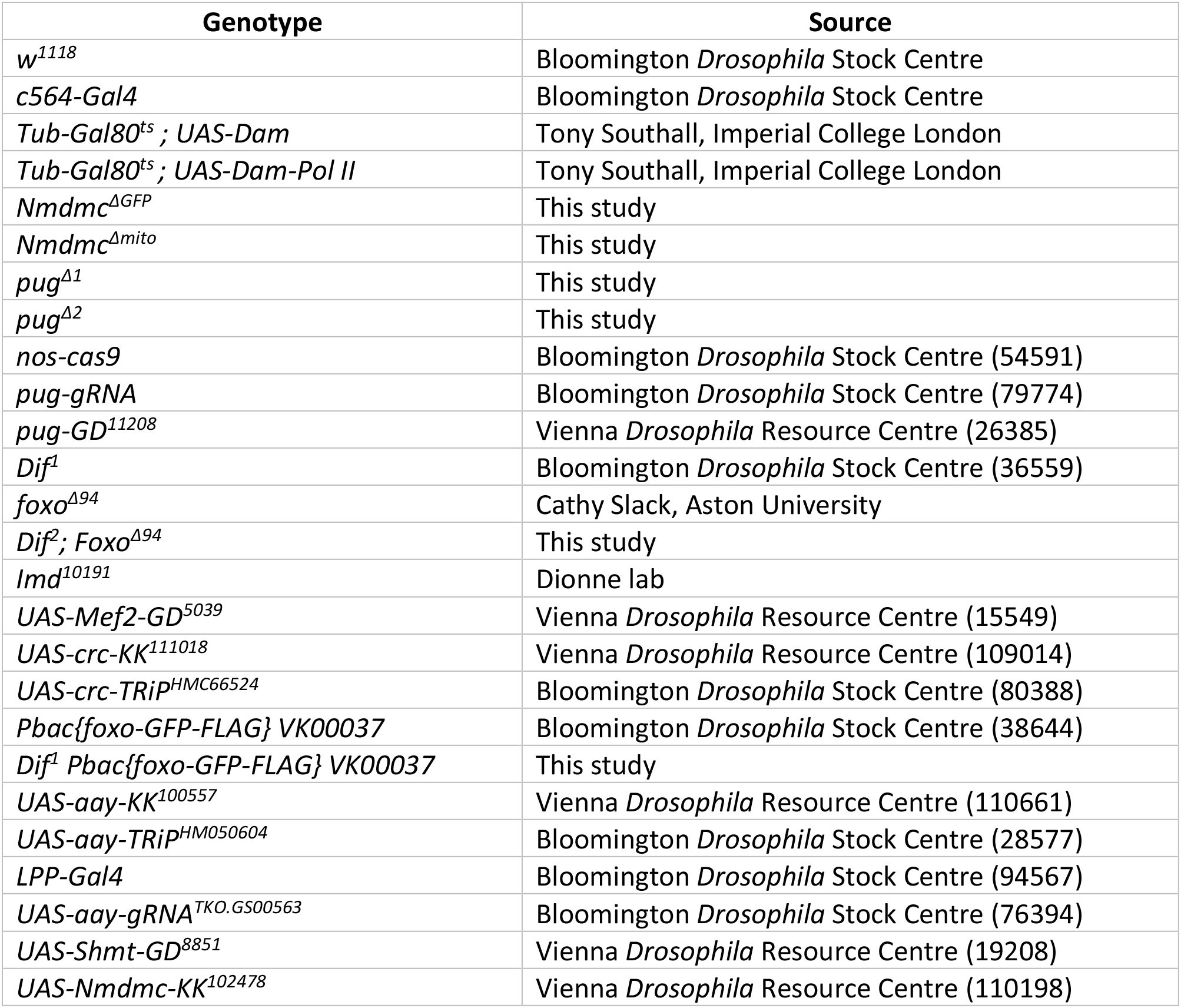
*Drosophila* lines used in this study.

### Bacterial infections

For all experiments, flies were collected within 2 days of eclosure and maintained on fresh food in groups of 20 for 5-7 days at 25 °C (unless otherwise stated). Bacteria were grown overnight at 37°C with shaking in the relevant media presented in Table 2. The following morning bacteria were pelleted and resuspended to the required optical density at 600 nm (OD600) as indicated in Table 2 (instances where different optical densities were used are noted in figure legends). 50 nL of bacterial suspension was injected into the fly abdomen using a Picospritzer III system (Parker Hannifin) and pulled glass capillary needles. As a wounding control, flies were injected with sterile PBS. As an anaesthesia control, flies were left on the CO2 pad for 5 mins but otherwise unmanipulated. Both experimental and control flies were then maintained at 29°C for the duration of the experiment.

**Table 2.**
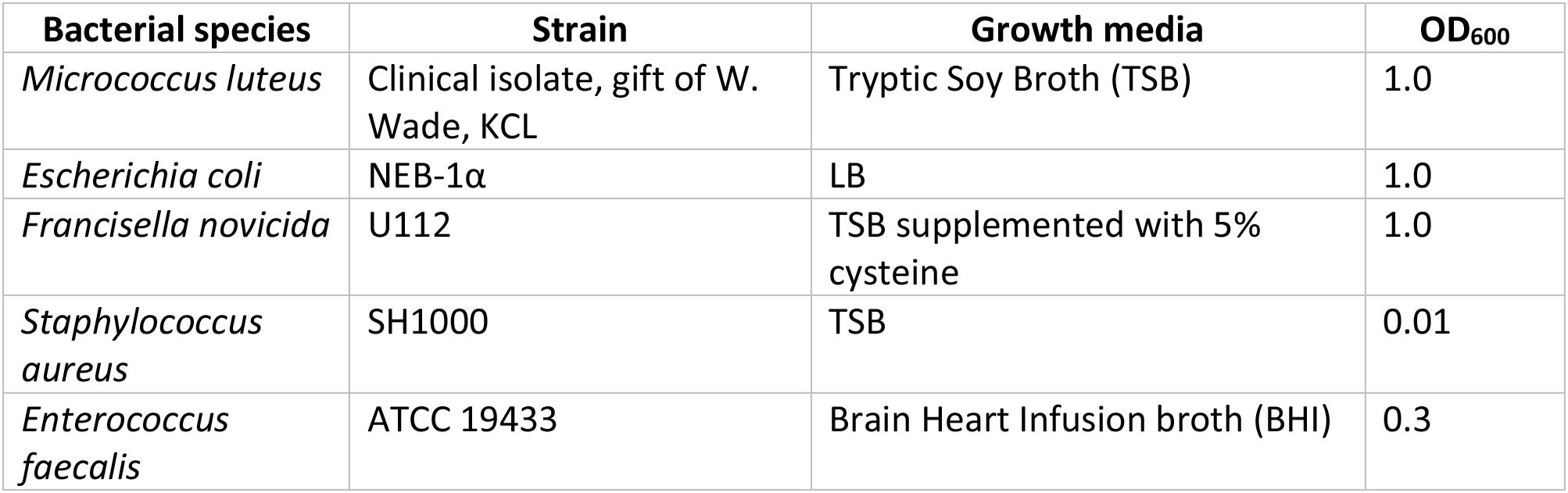
Bacterial strains, growth media and OD_600_ used for infections.

### RNAseq

*w^1118^*; *c564-Gal4* / + or *w^1118^*; *c564-Gal4* / +; *UAS-Mef2-IR* / + flies were smashed in Trizol (Invitrogen) and total RNA was extracted according to the manufacturer’s instructions. Library preparation and sequencing were carried out by UCL Genomics. Libraries were prepared using Illumina TruSeq Stranded mRNA v2, with the following changes from the manufacturer’s protocol: starting samples were 250ng total RNA; fragmentation was performed for 10 minutes at 94°C; no spike-ins were used; and 14 cycles of PCR were used to enrich libraries. At least 35 million reads per library were sequenced on an Illumina NextSeq 500 using v1 chemistry. Reads were assigned to transcripts using Kallisto^22^. Gene-level estimates for transcripts per million were extracted and differentially-expressed genes were identified using Sleuth^23^. Full RNAseq data are available at NCBI GEO as accession GSE216042.

### Targeted DamID

*c564-Gal4* flies were crossed to either *tub-Gal80^ts^; UAS-Dam* or *tub-Gal80^ts^; UAS-Dam-Pol II* flies at 18°C. Progeny were collected and maintained at 18°C for 7-10 days. Flies were shifted to 29°C to induce Dam expression, and infected 8 hours later. 20 flies per sample were flash frozen on dry ice 16 hours post-infection (*M. luteus*) or 66 hours post-infection (*F. novicida*). Frozen flies were vortexed and sieved to isolate the abdomen and thorax. Tissue was then homogenised in PBS supplemented with 83 mM EDTA. Once three independent replicates were obtained for each condition, samples were processed synchronously. Genomic DNA was extracted using a DNA micro kit (Qiagen), and then amplification of methylated DNA and preparation of sequencing libraries were performed as previously described^24,25^. Libraries were sequenced with Illumina Hiseq 50 bp single-end read sequencing.

### Targeted DamID analysis

Sequenced Dam-Pol II reads were normalised to Dam-only reads and mapped back to release 6.03 of the *Drosophila* genome using an previously described pipeline^26^. Genes showing significant Pol II occupancy were identified using a Perl script described in Mundorf et al. (2019)^27^ based on one developed by Southall et al. (2013)^24^ (available at-https://github.com/tonysouthall/Dam-RNA_POLII_analysis). *Drosophila* genome annotation release 6.11 was used, with 1% FDR and 0.3 log2 ratio thresholds. To identify genes that showed significantly different Pol II occupancy across treatment conditions, the log2(Pol/Dam) files from one treatment group were subtracted from another, producing three replicate comparison files. The same analysis script described above was run on the comparison files to identify genes with significantly different Pol II binding. Genes were filtered to eliminate those that did not show Pol II binding in the original data set.

### Quantitative real-time RT-PCR

Three flies per sample were smashed in TRI Reagent (Sigma) as per the manufacturer’s instructions. RNA was isolated via chloroform extraction and isopropanol precipitation. After washing in 70 % ethanol, the pellet was resuspended and treated with RNAse-free DNAse I (Thermo Fisher). The resulting RNA was then primed with random hexamers and reverse transcribed using RevertAid M-MuLV reverse transcriptase (Thermo Fisher). 10 μl of each sample was pooled and then serially diluted to generate a set of standards that were later used to create a standard curve in each experiment. The remaining 10 μl was diluted 1:40 and used for analysis.

qPCR was performed on a Corbett Rotor-Gene 6000 using SyGreen 2x qPCR mix (PCR Biosystems) with the primers indicated in Table S1. Cycling conditions were as follows: hold at 95°C for 10 minutes, then 45 cycles of 95°C for 15 s, 57°C for 30 s, and 72°C for 30 s, followed by a melting curve. Expression was determined by comparison of each value to the standard curve, followed by normalisation to housekeeping gene *RpL4.*

### CRISPR mutagenesis

To generate a mitochondrial *Nmdmc* mutant (*Nmdmc^Δmito^*), a guide RNA complementary to a portion of the predicted mitochondrial targeting sequence was designed and cloned into a pCFD5 backbone. The plasmid was injected into embryos expressing Cas9 (Cambridge Fly Facility), and stocks of potential mutants were created using a balancer line. Stocks were then screened for mutations via PCR amplification and Sanger sequencing of a 600 bp region of the *Nmdmc* sequence inclusive of the gRNA target site.

To generate a mutant lacking all isoforms of *Nmdmc* and expressing GFP in its place (*Nmdmc^ΔGFP^)*, we generated two guide RNAs complementary to the most 5’ exon common to all isoforms, targeting sites ~ 400 bp apart. These were cloned together into a pCFD5 backbone^28^. A second plasmid was generated in pScarlessHD -dsRed, in which in-frame GFP was inserted replacing most of the primary the *Nmdmc* coding exon, along with a 3xP3-dsRed cassette flanked by piggyBac transposable element ends^29^. This was flanked by ~ 1kb of sequence complementary to the region either side of the predicted break sites. This enables homologous recombination between the chromosome and the plasmid at the break sites, resulting in insertion of GFP and dsRed in place of the portion of the *Nmdmc* exon that is deleted in between the two gRNA target sites. These two plasmids were injected into embryos expressing Cas9 (Cambridge Fly Facility), and stocks of potential mutants were generated using a balancer line. Stocks were screened for mutations by the presence of 3xP3-dsRed, which was later flipped out using a piggybac transposase. Mutations were confirmed by PCR using primers that span the *Nmdmc-GFP* fusion junction. GFP expression was confirmed by western blot.

A fly line harbouring a gRNA targeting the most 5’ exon common to all *pug* isoforms was already available for use. Flies expressing *nos-Cas9* were crossed to flies expressing the *pug-gRNA*, and stocks were made from the potentially mutant offspring. Stocks were then screened for mutations via PCR amplification and Sanger sequencing of a 650 bp region of the *pug* sequence inclusive of the gRNA target site.

For all mutant lines, gRNAs and PCR primers are shown in Table S2.

### PCR

A single fly was homogenised in a solution of 10 mM Tris-Cl pH 8.2, 1 mM EDTA, 25 mM NaCl and 200 μg/ml Proteinase K (NEB, USA). The homogenate was heated at 37°C for 30 minutes followed by Proteinase K inactivation at 95 °C for 10 mins. PCR was performed using the primers listed in Table S2 and the following cycling conditions: hold at 94°C for 1.5 minutes, then 35 cycles of 94 °C for 30 s, 55 °C for 40 s, and 72 °C for 45 s, followed by a hold at 72 °C for 2 mins. Amplified products were visualised on a 1% agarose gel, purified and sent for Sanger sequencing where appropriate.

### Binding site prediction

Analysis of binding-site overrepresentation and identification of individual putative binding sites was performed using CLOVER^30^. The binding-site matrices used were: human FOXO3, JASPAR MA0157.2; mouse Foxo1, JASPAR MA0480.1; and *Drosophila* NF-κB binding sites^31–33^. The background sequence was *Drosophila* chromosome arm 3R. Input DNA sequences were Flybase “Extended Gene Regions” for *astray* and *Nmdmc*, corresponding to the full genomic sequence from first identified transcriptional start to last identified transcriptional stop, plus 2 kb on each end.

### Western blot

For each sample, three flies were homogenised in 75 μl Laemmli buffer supplemented with dithiothreitol. 5 μl of the homogenate was run on an SDS-PAGE gel, and proteins were then transferred to a PVDF membrane. The membrane was blocked in 5% milk diluted in TBS-T (Tris-buffered saline supplemented with 0.1% Tween-20), washed, and then incubated overnight with primary antibody diluted in Tris-buffered saline supplemented with 5% bovine serum albumen. After further washing, membranes were probed with secondary antibody for 1-2 hours, then imaged using SuperSignal West Pico PLUS Chemiluminescent substrate (Thermo Fisher) and on a ChemiDocTM XRS+ imaging system (Bio-Rad). Membranes were stripped for 1 hour in 0.2 M NaOH to enable re-probing. Primary antibodies and dilutions: anti-phospho-Akt (CST, 1:1000), anti-Akt (CST, 1:1000), anti-GFP (CST, 1:1000), anti-tubulin (DSHB, 1:5000). Secondary antibodies: horseradish peroxidase-conjugated anti-rabbit IgG and anti-mouse IgG (CST, 1:5000). Bands were quantified using ImageJ.

### Imaging

Abdominal fat body tissue was dissected from 5-8 day-old FOXO-GFP-FLAG-expressing flies and fixed at room temperature for 40 minutes in PBS with 4% paraformaldehyde. Preparations were incubated in PBS, 4% horse serum, 0.3% Triton X-100 at 4°C to block and then with anti-FLAG antibody (Sigma F1804, 1:1000) in the horse serum solution overnight. Subsequent washes were done at room temperature following standard protocols. Secondary antibody incubations were done at room temperature for 90 minutes. Cy3 - conjugated secondary antibodies were obtained from Jackson Immunoresearch and used at 1:100. Finally, preparations were stained with HCS LipidTOX Green (1:100, ThermoFisher Scientific) and DAPI (300 nM, ThermoFisher Scientific) for 15 minutes before mounting in glycerol with propyl gallate. Imaging was done using a Zeiss LSM710 confocal microscope and a 100x/1.46 Plan Apochromat oil immersion lens. Images were analysed using Fiji. All laser gains and offsets were kept constant to allow direct comparison of fluorescence between groups.

For quantification of FOXO nuclear signal, all laser gains and offsets were kept constant to allow direct comparison of fluorescence between groups. Briefly, images were visualised in the DAPI channel and nuclear regions of interest were selected and saved. The mean signal over these nuclear regions was then recorded in the FOXO channel and background subtracted. The FOXO signal over the entire fat body region was then recorded, background subtracted, and the ratio of nuclear to entire signal calculated. Three independent regions for each fat body (n=10 for each condition) were imaged. Images were analysed using Fiji.

### Metabolomics

For each sample, three flies were collected in cold LCMS solution, comprised of 2:2:1 methanol: acetonitrile: ultrapure water (all LCMS-grade). Flies were homogenised and stored at −80°C overnight or longer. After thawing at −20°C, samples were centrifuged at 15,000 x G at 4°C for 5 minutes. The supernatant was transferred to Spin-X 0.22 μm centrifuge filters and centrifuged at 15,000 x G at 4 °C for 75 minutes. 5 μl of filtrate from each sample was pooled, then the pool was serially diluted 1 in 2 to create a standard curve. An aliquot of each experimental filtered sample was diluted 1 in10 in LCMS solution as was an aliquot of the pool to create run controls. Immediately prior to analysis, the experimental samples, serially diluted pools, and run controls were then diluted 1:1 in acetonitrile containing 0.2% acetic acid. Samples were centrifuged at 17,000 x G at 4°C for 10 minutes and 100 μl of the supernatant taken for LC/MS analysis.

The supernatant was then transferred into LC/MS V-shaped vials (Agilent 5188-2788), and a 4 μl aliquot was injected into the LC/MS instrument. Aqueous normal-phase liquid chromatography was performed using an Agilent 1290 Infinity II LC system equipped with a binary pump, temperature-controlled autosampler (set at 4 °C) and temperature-controlled column compartment (set at 25 °C) containing a Cogent Diamond Hydride Type C silica column (150 mm × 2.1 mm; dead volume of 315 μl). A flow rate of 0.4 mL/min was used. The elution of polar metabolites was performed using solvent A, which consisted of deionized water (resistivity ~18 M⍰ cm) and 0.2% acetic acid, and solvent B, which consisted of 0.2% acetic acid in acetonitrile. The following gradient was applied at a flow rate of 0.4 ml/min: 0 minutes, 85% B; 0-2 minutes, 85% B; 3-5 minutes, 80% B; 6-7 minutes, 75% B; 8-9 minutes, 70% B; 10-11 minutes, 50% B; 11.1-14 minutes, 20% B; 14.1-25 minutes, 20% B; and 5-minutes of re-equilibration at 85% B. Accurate mass spectrometry was performed using an Agilent Accurate Mass 6545 QTOF apparatus. Dynamic mass axis calibration was achieved by continuous infusion after the chromatography of a reference mass solution using an isocratic pump connected to an ESI ionization source operated in positive-ion mode. The nozzle and fragmentor voltages were set to 2,000 V and 100 V, respectively. The nebulizer pressure was set to 50 psig, and the nitrogen drying gas flow rate was set to 5 l/minute. The drying gas temperature was maintained at 300 °C. The MS acquisition rate was 1.5 spectra/sec, and m/z data ranging from 50 to 1,200 were stored. This instrument enabled accurate mass spectral measurements with an error of less than 5 parts per million (ppm), a mass resolution ranging from 10,000 to 45,000 over the m/z range of 121-955 atomic mass units, and a 100,000-fold dynamic range with picomolar sensitivity. The data were collected in centroid 4-GHz (extended dynamic range) mode. The detected m/z data were deemed to represent metabolites, which were identified based on unique accurate mass-retention times and MS/MS fragmentation identifiers for masses exhibiting the expected distribution of accompanying isotopomers. The typical variation in the abundance of most of the metabolites remained between 5 and 10% under these experimental conditions.

### Metabolomics analysis

Mass spectrometry data was analysed using the Agilent Masshunter software suite, Microsoft Excel and in-house R scripts.

Features were identified as the compounds of interest by comparison with metabolite standards. To quantify metabolite abundances, peak heights and areas were extracted using Agilent Masshunter Profinder. Features identified as serine, glycine, and threonine were quantified in addition to the most abundant 100 features. Once abundances were corrected by subtraction of any background signal found in blank control runs, quality control was performed by examination of the pooled-sample serial dilution and run controls. Features with an r^2^ < 0.85 in the serially diluted samples, or a relative standard deviation of > 0.25 in the run controls were removed from analysis. The abundances of the metabolites of interest were then normalised to the sum of the abundances of all quantified features in each sample.

### Survival assays

For some experiments, survivals were monitored manually. Flies were housed in groups of 20 at 29°C and deaths were monitored daily. Flies were transferred onto fresh food every three to four days.

For other experiments, survival was monitored using ethoscopes^34^. These devices allow constant monitoring of fly survival using video tracking software. After infection, flies were sorted into individual glass tubes (70 mm x 5 mm x 3 mm (length x external diameter x internal diameter)) containing standard food and loaded into ethoscopes. Survival was monitored for 4-5 days at 29 °C and 60 % humidity. Survival was monitored for 4-5 days at 29 °C and 60 % humidity. We defined the time of death as the last detected movement before a 6h period of constant immobility. Analysis was performed in RStudio using the Rethomics suite of packages^35^.

For both approaches, survivals were visualised in RStudio using the Kaplan-Meier method, and survival differences between infected flies were compared statistically using a log-rank test.

### Bacterial quantifications

Individual flies were homogenized in 100 μl sterile PBS. Homogenates were serially diluted and plated onto either TSB or BHI agar plates, for *S. aureus* and *E. faecalis* respectively. Plates were incubated at room temperature overnight, and then placed at 37°C the following morning until colonies were large enough to count. Back calculations were performed on the colony counts to determine the number of colony-forming units (CFU) originally present in each fly.

### Graphing and Statistical analysis

Graphing and statistical analysis were performed in R (version 4.1.2) using RStudio (build 492). For all data except survivals, a Shapiro-Wilk test was first performed to test normality. For pairwise comparisons, either a t-test or a Wilcoxon test was performed, for normally and non-normally distributed data respectively. For multiple comparisons, ANOVAs were performed. For grouped data showing a normal distribution, comparisons were made using a one-way ANOVA with a post-hoc Tukey test. For non-normal grouped data, a Kruskal Wallis ANOVA was performed with a post-hoc Dunn test, and Benjamini-Hochberg correction for multiple comparisons.

## Results

### Serine, glycine and folate metabolism is transcriptionally regulated during infection

To identify novel genes that may play important roles in the immune response, we performed RNAseq on infected flies. Flies were infected with *Micrococcus luteus* or *Escherichia coli*, non-lethal bacteria that evoke a strong immune response. Gene expression was compared to that of uninjected flies, or flies given a sterile PBS injury, three hours post-infection (Tables S3 and S4).

We observed strong transcriptional regulation of *astray* (*aay*), *Shmt*, *Nmdmc*, and *pugilist* (*pug*)(Fig 1A). These genes encode metabolic enzymes with closely connected functions (Fig 1B). *astray* encodes phosphoserine phosphatase, which converts free phosphoserine generated from glycolytic intermediates into serine. This serine can be converted into glycine. The interconversion of serine and glycine, accomplished by serine hydroxymethyltransferase (Shmt), is a critical interaction point between amino acid metabolism and the folic acid cycle, simultaneously interconverting tetrahydrofolate (THF) and 5,10-methylene-THF. *Nmdmc* encodes NAD-dependent methylenetetrahydrofolate dehydrogenase, a bifunctional enzyme that interconverts 5,10-methylene THF and 10-formyl-THF, simultaneously interconverting NAD+ and NADH. *pug* encodes an enzyme with similar activity, but dependent on NADP+ and NADPH. The module encoded by *astray*, *Shmt*, *pug*, and *Nmdmc* thus is a nodal point connecting amino acid metabolism, one-carbon metabolism, redox state, and glycolysis. The regulation of *astray* and *Nmdmc* was particularly striking, as these enzymes were among the few metabolic genes that were transcriptionally induced, as opposed to repressed, by infection (Fig 1A, Table S3). The upregulation of *astray* and *Nmdmc* during *M. luteus* infection was confirmed by qPCR (Fig S1A). Given their potential biological significance during infection, we decided to further investigate the regulation and role of these enzymes.

**Figure 1.**
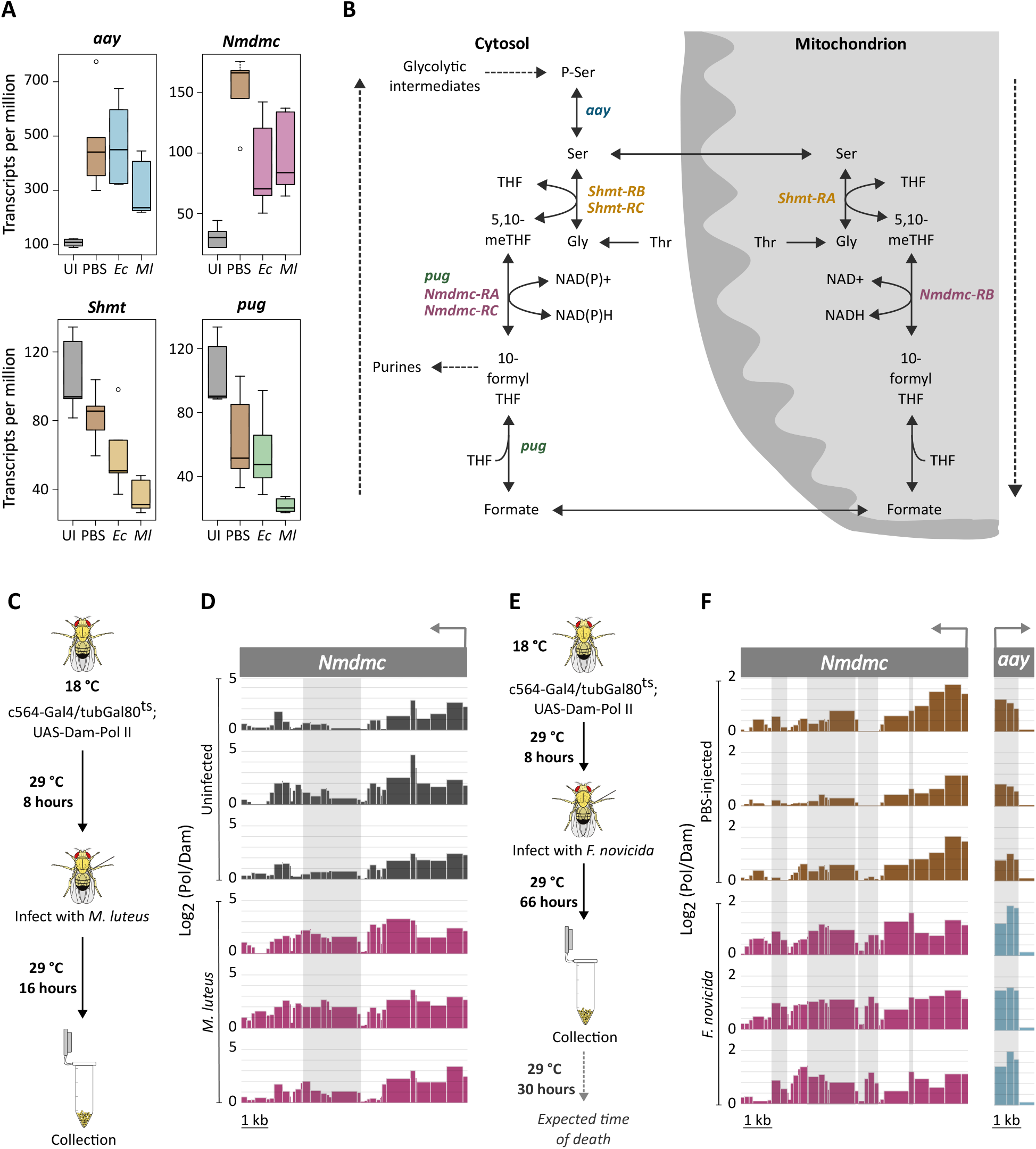
Transcriptional regulation of serine, glycine, and folate metabolism during infection. **A.** Expression of *astray*, *Nmdmc*, *Shmt*, and *pug* (D) 3h after infection with *E. coli* (*Ec*) or *M. luteus* (*Ml*). Measurements made by RNA-seq, n=5. **B.** Schematic of serine, glycine, and folate metabolism. **C, E**. *M. luteus* (C) and *F. novicida* (E) TaDa protocol schematics. **D, F**. Pol II binding at *Nmdmc* and *astray* during *M. luteus* (D) and *F. novicida* (F) infection. Y axis indicates log2 ratio between Dam-Pol II and Dam-only mapped reads. Light grey bars indicate regions of increased binding in infected samples. Each track represents an independent repeat.

### *astray* and *Nmdmc* are upregulated in the fat body during infection

The fat body plays important immune and metabolic roles, so we sought a readout of fat body-specific gene expression during infection. To achieve this, we performed Targeted DamID (TaDa)^24^. A subunit of RNA polymerase II fused to a bacterial DNA adenine methylase (Dam) was expressed specifically in the fat body using the *c564-Gal4* driver. *Drosophila* does not exhibit endogenous adenine methylation; by identifying sites of Dam-mediated adenine methylation, it is therefore possible to identify locations of RNA polymerase binding and thus genes that are actively transcribed.

To prevent methylation during development, we utilised the temperature sensitive Gal80 repressor and reared the flies at 18°C. Eight hours prior to infection, flies were temperature shifted to 29°C to induce Dam-Pol II expression. To ensure sufficient time was left for detectable methylation markings to be deposited by Dam, flies were collected 16 hours after injection of *M. luteus* (Fig 1C). The PBS-injection control binding profiles looked broadly similar to the *M. luteus* profiles, likely due to the PBS injection evoking a transient immune response. Hence comparisons were made between the infected samples and uninfected controls. The full list of genes upregulated upon infection is shown in Table S5. Significantly increased Pol II binding was observed at the *Nmdmc* locus (Fig 1D). Because adenine methylation is very stable in *Drosophila*, reliable identification of genes downregulated by infection was not possible.

Though *M. luteus* evokes a strong immune response, we were concerned that its rapid rate of clearance would mean that transient gene expression changes were not detectable by TaDa. We were unable to identify a Toll pathway agonist that would drive persistent, strong immune activation without killing the fly, so instead used *Francisella novicida*. This is a Gram-negative bacterium that causes a chronic infection, resulting in highly synchronous death several days after infection^36^. We confirmed that *astray* and *Nmdmc* are upregulated in the whole fly during *F. novicida* infection by qPCR (Fig S1B), and then performed TaDa with a 66-hour collection timepoint (Fig 1E). The full list of upregulated genes is presented in Table S6. Both *astray* and *Nmdmc* showed significantly increased Pol II binding upon *F. novicida* infection (Fig 1F), confirming that infection-induced upregulation of both genes occurs in the fat body.

### Transcriptional preference is shown towards mitochondrial isoforms during infection

We next investigated how expression of the individual isoforms of the enzymes involved in serine, glycine and 1C metabolism were impacted by infection. Only one isoform of *astray* exists (Fig 1B). Five isoforms of *pug* exist, all of which are predicted to be cytosolic. *Nmdmc* and *Shmt* each have three isoforms, two of which are predicted to be cytosolic, and one mitochondrial. Bar *astray*, each of these enzymes are expected to act reversibly. However, evidence from mammalian cell culture, animal models and human flux studies show that it is usually thermodynamically favourable for the reactions to run oxidatively in the mitochondria in the direction of serine to glycine conversion, and in the opposite direction in the cytosol^37,38^ (indicated by the dotted clockwise arrows in Fig 1B). We sought to determine whether there was any difference in transcriptional regulation of the cytosolic and mitochondrial isoforms during infection. We focussed on *Staphylococcus aureus* for the remainder of the study as this is a Toll-pathway agonist that would not only be an immune elicitor but also a possible driver of pathology.

As with the other infections tested, increased expression of *astray* was observed six hours post-infection with *S. aureus* (Fig 2A). Increased expression of *Nmdmc* was also observed, but upon studying the individual isoforms, we found that the upregulation was limited to the mitochondrial isoform, *Nmdmc-RB* (Fig 2B). Again, in accordance with the other infections studied, we observed reduced expression of *Shmt* upon *S. aureus* infection (Fig 2C). This reduction, however, was limited to the cytosolic isoforms of *Shmt*, with the expression of mitochondrial *Shmt* remaining unchanged (Fig 2C). We again observed a reduction in *pug* expression (Fig 2D), which is purely cytosolic. Together this indicates that there is a clear difference in the regulation of these metabolic enzymes based on subcellular localisation. Mitochondrial isoforms are either unchanged or upregulated upon infection, whilst cytosolic isoforms are unchanged or downregulated. Hence there is a transcriptional preference towards the mitochondrial pathway during infection.

**Figure 2.**
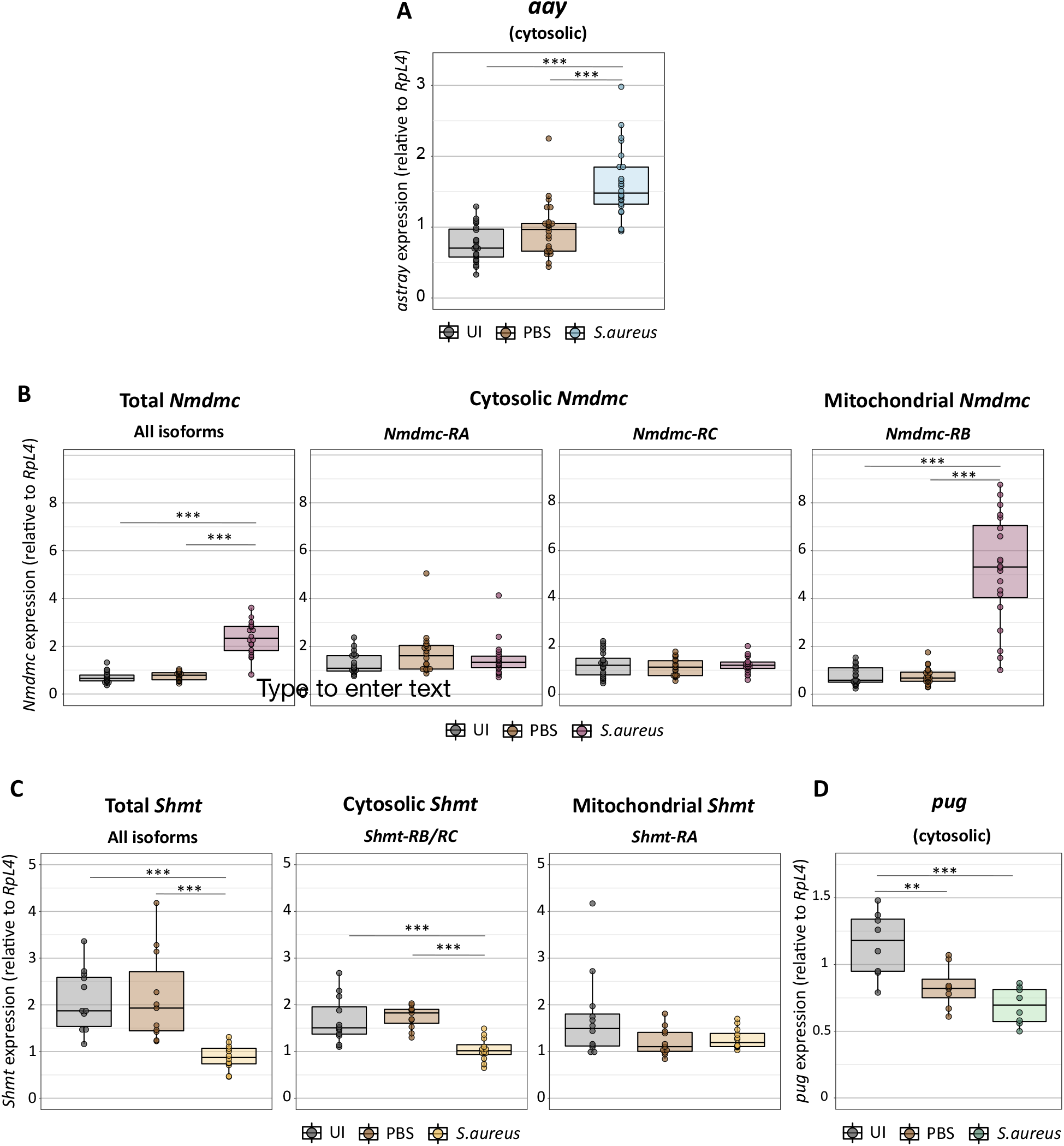
Isoform-specific regulation of serine-folate metabolic unit enzymes upon infection. **A-D**. Expression of *astray* (A), *Nmdmc* (B), *Shmt* (C), and *pug* (D) isoforms in c564-Gal4>+ flies 6h after *S. aureus* infection. Measurements made by RT qPCR and normalised to expression of *RpL4*. A: n=24; B: n=20; C: n=12; D: n=8. * p<0.05; ** p<0.01; *** p<0.001. Lack of stars indicates no significant difference. Statistical comparisons were done using Kruskal-Wallis ANOVA (A-C) or parametric ANOVA (D), depending on normality of underlying data. UI = uninfected.

### *Nmdmc* and *pug* show transcriptional compensation

In mammalian cell culture, it has been shown that deletion of the homologue of *Nmdmc* results in compensation from the cytosolic pathway^38^. To determine whether the same compensatory effect occurs in *Drosophila*, we created *Nmdmc* CRISPR mutants. We generated a full *Nmdmc* mutant lacking all three isoforms, but found that these flies had poor viability, precluding adult infection experiments. In addition, we targeted solely the predicted mitochondrial isoform of *Nmdmc* (encoded by *Nmdmc-RB*) as this is upregulated during infection. We obtained viable mutant flies carrying a 12-nucleotide deletion that eliminated the start codon for this isoform; we named this allele *Nmdmc^Δmito^* (Fig 3A). We confirmed the lack of *Nmdmc-RB* expression by qPCR (Fig 3B). The coding sequence of *Relish*, the terminal transcription factor in the Imd signalling pathway, is located within an intron of *Nmdmc*, so we confirmed that *Relish* expression was unaffected by mitochondrial *Nmdmc* mutation (Fig S2A, B).

**Figure 3.**
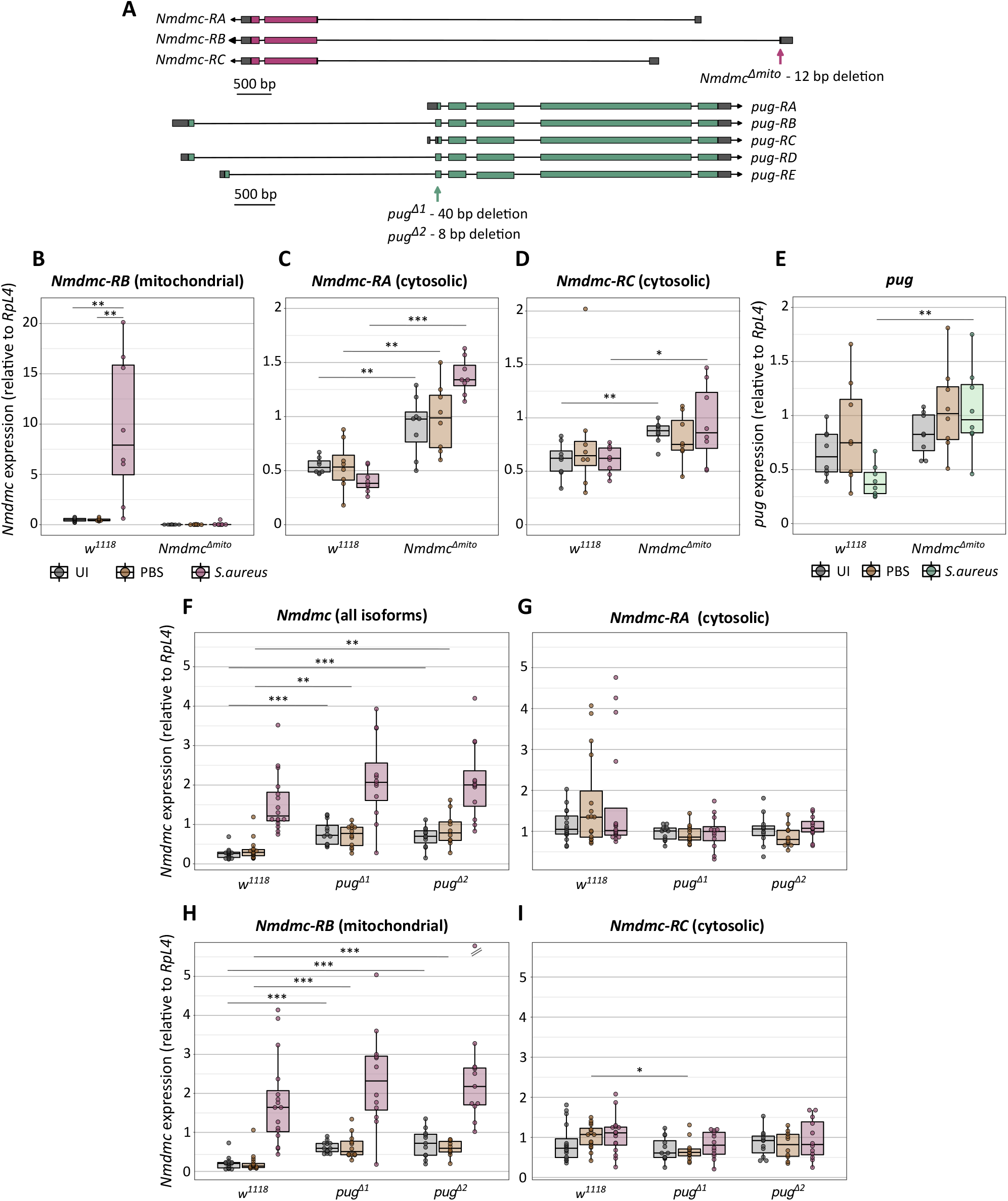
*Nmdmc* and *pug* show transcriptional compensation. **A**. Schematic showing *Nmdmc* and *pug* mutant alleles. **B-I**. Expression of *Nmdmc* isoforms and pug in Nmdmc mitochondrial mutants (B-E) and pug mutants (F-I), 6h after *S. aureus* infection. Measurements made by RT qPCR and normalised to expression of *RpL4*. n=8-20. B, stars indicate comparisons within genotypes; C-I, stars indicatges comparisons among genotypes. * p<0.05; ** p<0.01; *** p<0.001. Lack of stars indicates no significant difference. Statistical comparisons were done using Kruskal-Wallis ANOVA (B, F, G, I), t-test (C, E), or Wilcoxon test (D), depending on normality of underlying data. The out-of-range sample in H has a value of 12.2.

In healthy *Nmdmc^Δmito^* flies, we observed increased expression of both remaining (predicted cytosolic) isoforms of *Nmdmc* (Fig 3C-D). During infection, expression of both isoforms was further increased, particularly *Nmdmc-RA*. We observed a marginal increase in the expression of *pug* in healthy *Nmdmc^Δmito^* flies (Fig 3E). Further, we found that during infection *pug* was not downregulated in *Nmdmc^Δmito^* flies, as observed in wild-type controls, but maintained at a constant level (Fig 3E). Together, this indicates that expression of *pug* and the cytosolic isoforms of *Nmdmc* can increase to compensate for the lack of mitochondrial *Nmdmc*.

To determine whether this compensation is bidirectional, we generated *pug* mutants. *pug^Δ1^* and *pug^Δ2^* carried 40 bp and 8 bp deletions, respectively, in the most 5’ coding exon common to all *pug* isoforms (Fig 3A). Both mutations induced frame shifts that result in premature stop codons. During infection, expression of *Nmdmc* was not significantly changed in the *pug* mutants compared to the wild types (Fig 3F). However, in uninfected and PBS-injected *pug* mutants, we did observe increased *Nmdmc* expression (Fig 3F). Looking at the individual isoforms, we found that *Nmdmc-RB*, rather than the cytosolic isoforms, showed upregulation (Fig 3G-I). The same compensation from predicted mitochondrial *Nmdmc* was observed upon fat body knockdown of *pug* (Fig S3A-D). The fact that loss of a cytosolic enzyme is compensated for by a mitochondrial one, when alternative cytosolic equivalents are available, adds weight to the premise established in mammalian cell culture that the mitochondrial pathway is used preferentially^38^. This preference could be related to cofactor availability in the respective compartments, as *Nmdmc* is dependent on NAD whilst *pug* requires NADP (Fig 1B).

Together, these data demonstrate a bidirectional transcriptional compensation between *pug* and *Nmdmc*. Given that these enzymes are at least partially redundant in mammalian cell culture^38^, the observed compensation supports the hypothesis that they do perform their predicted functions based on homology to mammalian counterparts.

### *astray* and *Nmdmc* are upregulated by Dif and FOXO and repressed by MEF2

We next investigated the mechanisms by which *astray* and *Nmdmc* are upregulated during infection. To achieve this, we tested expression of *astray* and *Nmdmc* in flies mutant for *Dif* (the primary transcriptional mediator of Toll signalling in adult flies), *imd* (a critical adaptor in the Imd signalling pathway), and *foxo*, a major metabolic transcriptional regulator whose activity increases during infection^11^. In an attempt to find mechanisms consistent across infection, we infected flies with three different Gram-positive bacteria: *S. aureus, M. luteus* and *Enterococcus faecalis.*

*astray* and *Nmdmc* were still upregulated in *imd* mutants 6 hours post-infection; this was not unexpected, as the Imd pathway is primarily responsive to Gram-negative infection (Fig 4A-B). However, mutation of *Dif* eliminated the induction of *astray* in *M. luteus* and *E. faecalis* infection and reduced the magnitude of upregulation with *S. aureus* (Fig 4A). Similarly, *Nmdmc* upregulation was completely lost in *Dif* mutants (Fig 4B). Mutation of *foxo* had a similar effect, eliminating induction of both *astray* and *Nmdmc* (Fig 4A, B). The level of *astray* expression in uninfected and PBS-injected *foxo* mutants was considerably lower than in wild-type controls, suggesting that FOXO contributes to baseline *astray* expression in healthy flies, in keeping with the observation that the *astray* enhancer is bound by FOXO^39^ (Fig 4A). Given the dependence on both Dif and FOXO, we generated *Dif*-*foxo* double mutants, and again observed a complete absence of induction of *astray* and *Nmdmc* (Fig 4A-B). Together, this indicates that functional Toll signalling and FOXO activity are required for *astray* and *Nmdmc* upregulation during infection.

**Figure 4.**
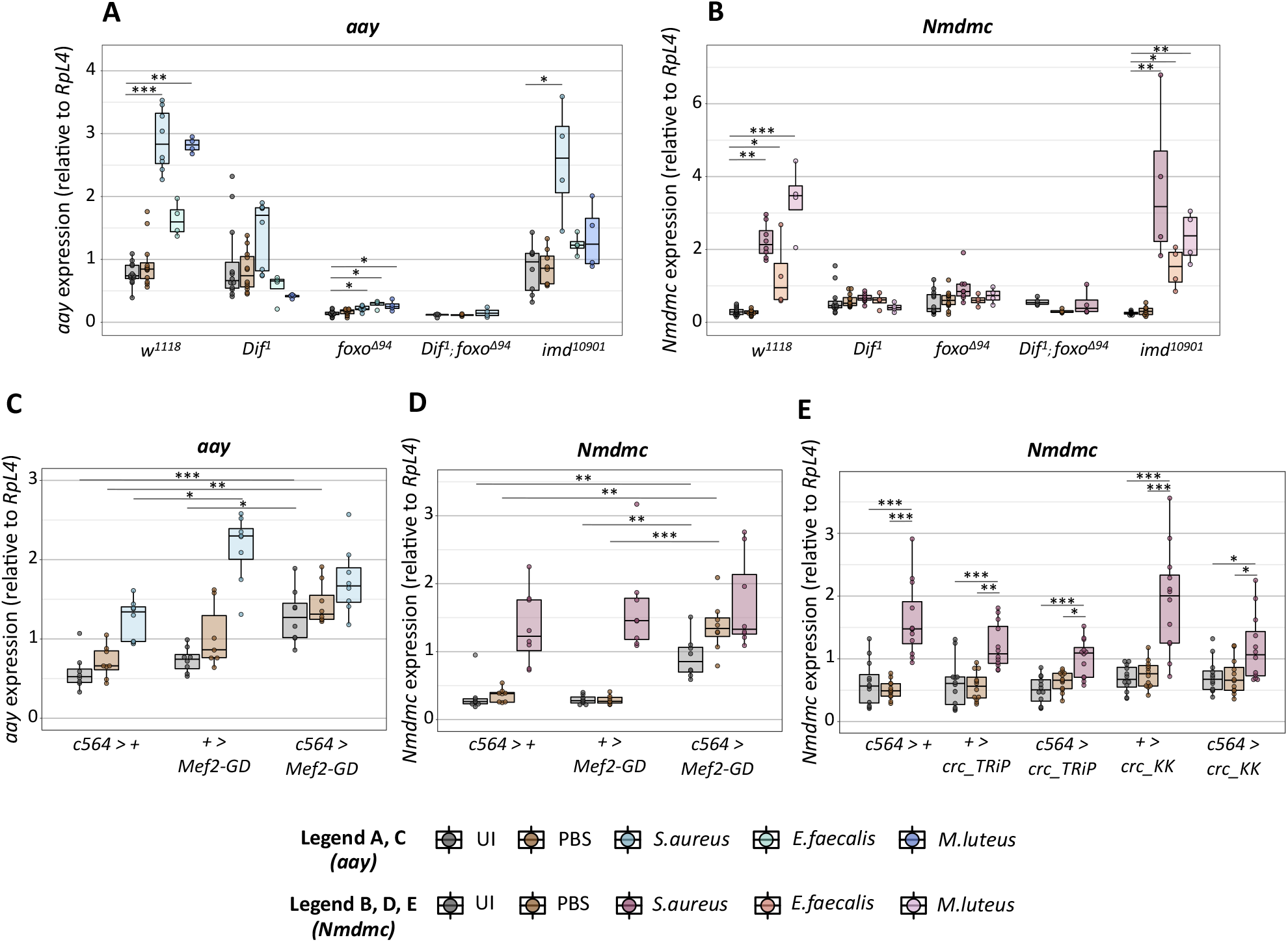
*Nmdmc* and *aay* are upregulated by Dif and FOXO and downregulated by MEF2. **A-B.** Expression of *astray* (A) and *Nmdmc* (B) in flies mutant for *Dif, foxo, imd*, or *Dif-foxo* double mutants 6h after infection with *S. aureus*, *E. faecalis*, or *M. luteus*. **C-D**. Expression of *astray* (C) and *Nmdmc* (D) in *Mef2* fat body knockdowns, 6h after infection with *S. aureus*. **E**. Expression of *Nmdmc* in two different fat body *crc* knockdown lines, 6h after infection with *S. aureus*. All measurements made by RT qPCR and normalised to expression of *RpL4*. n=8-12. * p<0.05; ** p<0.01; *** p<0.001. Lack of stars indicates no significant difference. Statistical comparisons were done using Kruskal-Wallis ANOVA. UI = uninfected.

We next investigated whether *astray* and *Nmdmc* could be regulated by MEF2, an infection-responsive transcription factor known to drive expression of both immune and metabolic genes^16^. Surprisingly, fat body knock down of *Mef2* did not block *astray* or *Nmdmc* induction upon *S. aureus* infection, but instead resulted in increased expression of both enzymes in the uninfected and PBS-injected controls (Fig4C-D). This suggests that MEF2 may act as a repressor of *astray* and *Nmdmc*, consistent with the fact that MEF2 is able to recruit histone deacetylases to repress gene expression in mammals^40–43^.

Finally, given the importance of astray and Nmdmc in amino acid synthesis, we hypothesised that another plausible regulator could be ATF4. This transcription factor is activated by low amino acid availability for translation to drive a gene expression program to restore amino acid homeostasis^44^. Knock down of *crc*, the gene encoding ATF4, in the adult fly brain has previously been shown to reduce *Nmdmc* expression^45^. Further, knock down of *ATF4* in mammalian cell culture has been shown to reduce expression of the homologues of both *astray* and *Nmdmc*^46^. We knocked down *crc* in the fat body using two independent RNAi lines (Fig S4A) and found an apparent reduction in the magnitude of *Nmdmc* induction upon infection (Fig 4E). However, the effect on *astray* was inconsistent, with *crc* knockdown reducing *astray* upregulation in some experiments but not others (Fig S4B-D). We did not observe any major differences in *Shmt* expression in *crc* knockdowns (Fig S4E), despite ATF4 having been shown to regulate S*hmt* in mammalian cell culture and the adult fly brain^45,46^.

Taken together, these data suggest that expression of *astray* and *Nmdmc* may be maintained at a low level in healthy flies by MEF2, but that upon infection the action of Dif, FOXO, and potentially ATF4 in the case of *Nmdmc*, is able to overcome the repression to drive upregulation.

### Dif and FOXO may act in a coordinated manner to drive *astray* and *Nmdmc* upregulation

Having identified Dif and FOXO as the primary transcription factors driving *astray* and *Nmdmc* induction, we sought to determine whether they were direct regulators or acting via an intermediate. *In silico* predictions indicated that FOXO binding sites were over-represented in the vicinity of *aay* and *Nmdmc* (p=0.012), while NF-κB sites showed no over-representation (p>0.1) (individual high-scoring FOXO sites on both loci are mapped in Fig 5A). FOXO is capable of physically binding the *astray* promoter *in vitro* and driving *astray* expression in S2 cells^39^, lending weight to the hypothesis that it could be a direct regulator of both genes *in vivo*.

**Figure 5.**
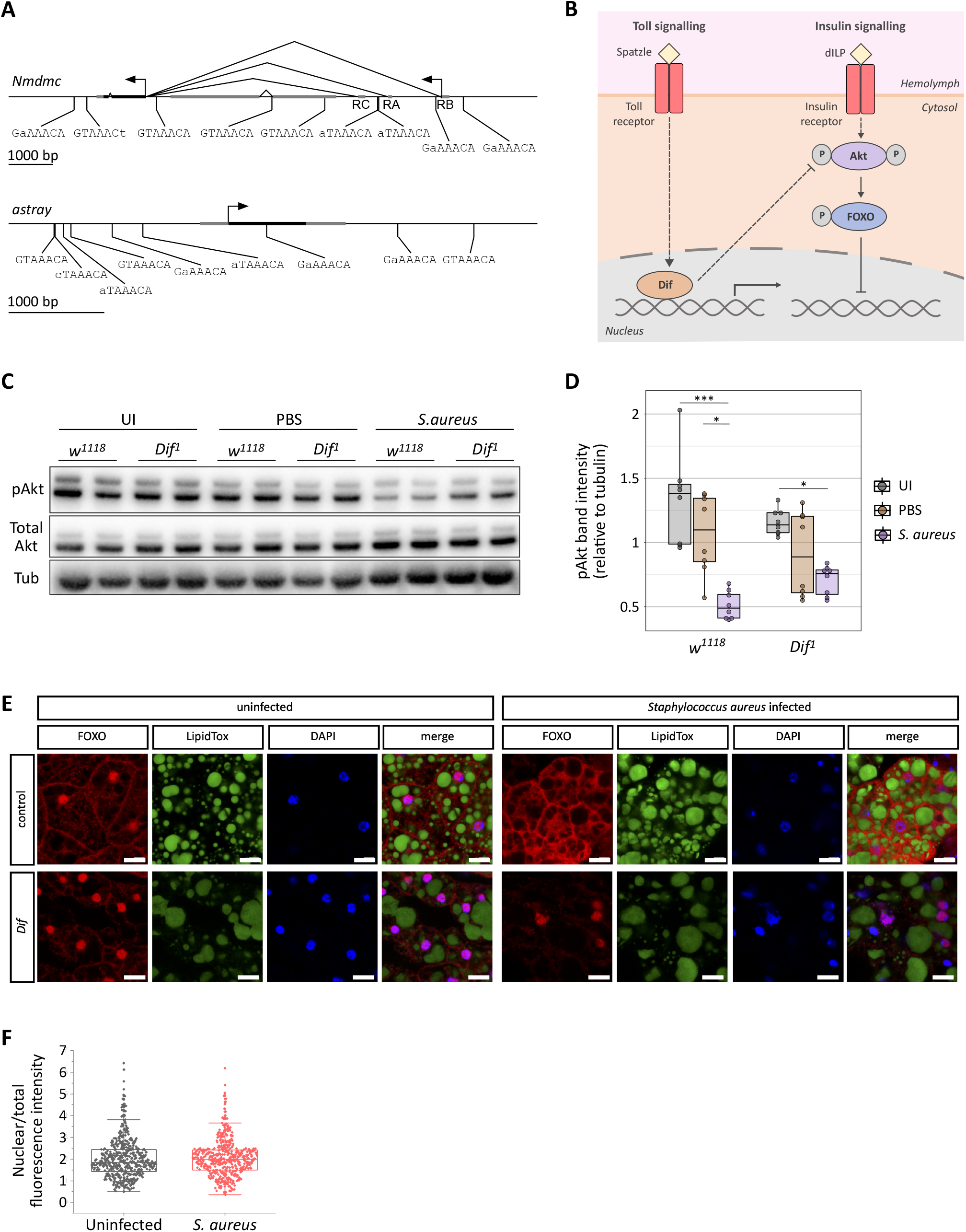
Dif and FOXO may act together to drive *astray* and *Nmdmc* upregulation. **A.** Schematic indicating predicted FOXO binding sites in the vicinity of *astray* and *Nmdmc*. Translational start sites are indicated by arrows and untranslated regions and other genes are shown in grey. Sites shown are either perfect matches to the 7mer FOXO consensus binding site (GTAAACA), or have one mismatch. **B.** Schematic of the interaction between Toll and insulin signalling. **C-D**. Western blot of pAkt, total Akt and tubulin in *w1118* and *Dif* flies 6h after *S. aureus* infection. C shows one blot representative of two independent repeats; D shows a quantification of the band intensities from both repeats. n = 8. In D, stars indicate significant differences between treatment groups within the same genotype.* p < 0.05, *** p < 0.001. Lack of stars indicates no significant difference. Statistical comparisons performed using Kruskal-Wallis ANOVA. UI: uninfected. **E**. FOXO, neutral lipid (LipidTox) and DAPI localisation in wild-type and *Dif* mutant fat body, uninfected and 6h after *Staphylococcus aureus* infection. Scale bar, 10 μm. **F**. Quantification of nuclear vs. total FOXO fluorescence intensity in fat body from *w1118* flies, uininfected or 6h after infection with *S. aureus*.

It is plausible that Dif could drive *astray* and *Nmdmc* induction via FOXO itself. FOXO is negatively regulated by the insulin signalling pathway; the insulin-responsive kinase Akt phosphorylates FOXO, preventing its nuclear translocation (Fig 5B). Toll signalling can disrupt the insulin signalling pathway in a Dif-dependent manner^11^. Active Toll signalling can reduce phosphorylation of Akt, resulting in lower phosphorylation and therefore increased FOXO activity (Fig 5B). Hence, we hypothesised that the inhibition of insulin signalling via the Toll pathway could be the reason that *astray* and *Nmdmc* show Dif dependency.

To date, the reduction in phosphorylated Akt (pAkt) as a result of Toll signalling has only been documented during *M. marinum, M. luteus* and *B. bassiana* infection of adult flies^9,11^ and through constitutive activation of the Toll receptor in larval fat bodies^11,13^. Therefore, we first sought to determine whether *S. aureus* infection induced the same disruption of insulin signalling; we observed significantly reduced levels of pAkt six hours post-infection (Fig 5C-D). In *Dif* mutants, the magnitude of pAkt reduction was smaller than in controls (Fig 5C-D), demonstrating that the disruption of insulin signalling in *S. aureus* infection is Dif-dependent. Hence, the observed requirement of Dif in *astray* and *Nmdmc* induction could be due to its ability to promote FOXO activity.

To further explore this possibility, we imaged FOXO localisation in adult fat bodies six hours post infection with *S. aureus*, in wild-type flies and *Dif* mutants (Fig 5E). In keeping with previous observations on adult *Drosophila* fat body, we found that FOXO localisation was highly variable but, in general, mostly nuclear in samples from wild-type flies that had not been infected^47,48^. This subcellular localisation was unchanged after infection of wild-type flies, suggesting that the regulation of FOXO activity in adult *Drosophila* fat body is different from what has previously been reported in other cell types (Fig 5F).

### Knockdown of *astray* or *Shmt* significantly perturbs serine, glycine and threonine metabolism

Having explored the transcriptional regulation of the enzymes in the serine-folate metabolic unit, we next investigated their functional relevance. To achieve this, we performed liquid chromatography mass spectrometry (LC/MS) on *astray* and *Shmt* fat body knock downs. As shown in Figure 6A, serine can be converted to glycine, which can also be derived from threonine. In keeping with our transcriptional data, serine abundance was increased by infection in control genotypes relative to PBS-injected controls (Figure 6B, C). The levels of serine, glycine, and threonine were markedly perturbed by *astray* and *Shmt* fat body knock down (Figure 6B, C). In *astray* knock downs, serine, glycine and threonine levels were all significantly lower than the genetic controls, at both six and 24 hours post-infection (Figure 6B). In *Shmt* knock downs, glycine and threonine were significantly reduced, but serine levels were significantly increased (Figure 6C). This suggests that *Shmt* predominantly runs in the direction of serine to glycine conversion, as loss of *Shmt* results in a build-up of serine at the expense of downstream glycine. The reduction in threonine is likely indicative of glycine being derived from threonine in the absence of serine.

**Figure 6.**
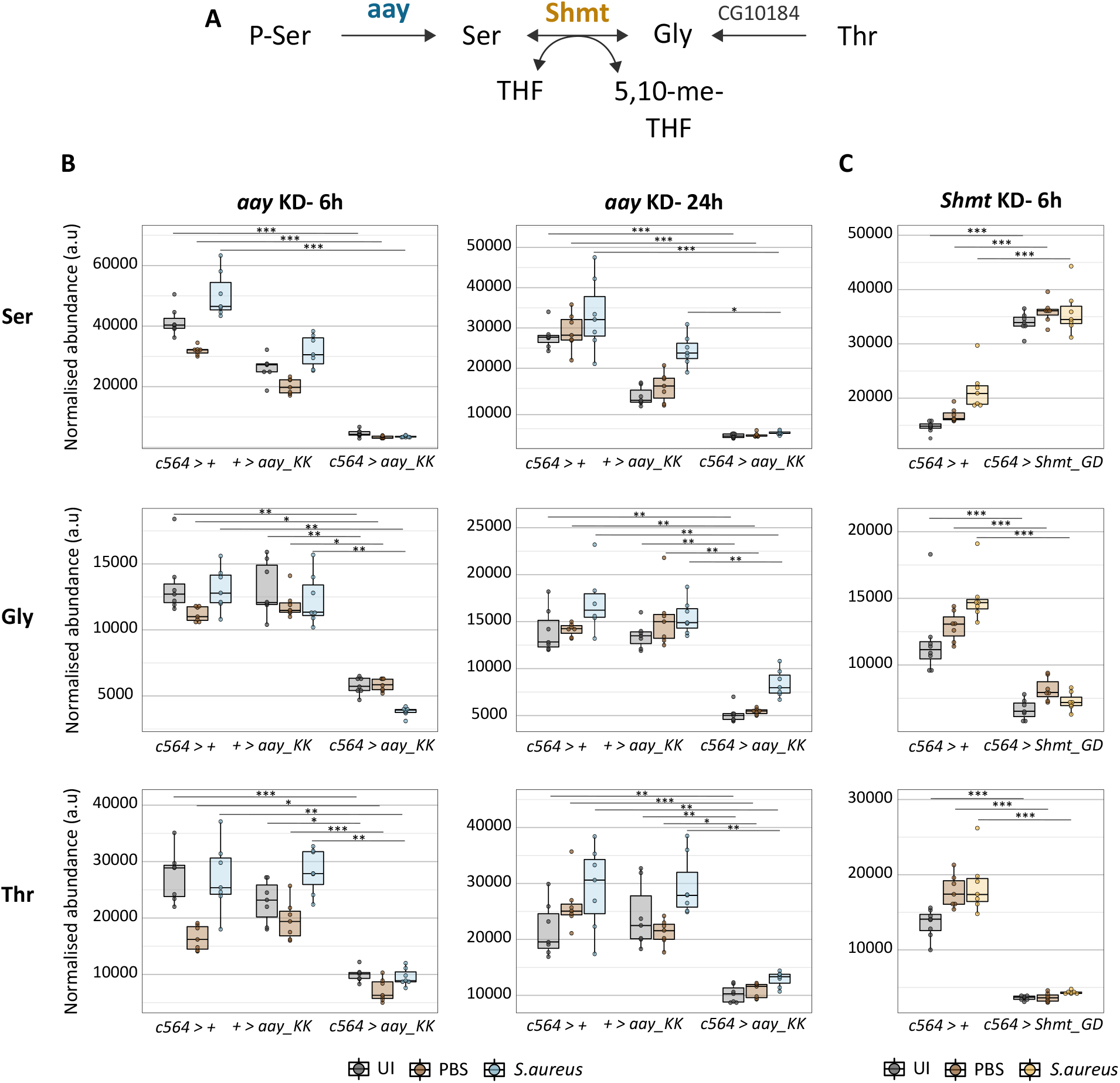
Metabolomic analysis of *astray* and *Shmt* fat body knockdowns. **A**. Schematic showing the relationship between serine, glycine, threonine. **B-C**. Levels of serine, glycine, and threonine in *astray* (B) and *Shmt* (C) fat body knockdowns, 6 or 24 hours after *S. aureus* infection. Assayed by LC-MS. n=7-8. * p<0.05; ** p<0.01; *** p<0.001. Lack of stars indicates no significant difference. Statistical comparisons for *astray* knockdowns were done using Kruskal-Wallis ANOVA, *Shmt* knockdown using Wilcoxon test, except for threonine, where normally-distributed data allowed use of a parametric t-test.

Together, these findings provide strong evidence that astray and Shmt perform their predicted enzymatic functions in *Drosophila melanogaster*. The fact that the metabolic perturbations resulted from fat body knock down of *astray* and *Shmt* demonstrates that the expression of these enzymes in the fat body specifically is important for whole-organism metabolism.

### Loss of *astray* and *foxo* expression impairs immune competence

Looking next at immune function, we observed that *astray* fat body knockdowns had a significant survival defect during *S. aureus* infection (Fig 7A). The same phenotype was observed upon knockdown of *astray* using a different RNAi line (Fig S5B-C) and fat body mutation of *astray* using tissue-specific CRISPR (Fig S5D). No meaningful difference in bacterial load was observed in *astray* knockdowns, with as great a difference observed between the two genetic controls as between the controls and the knockdown (Fig 7B). This demonstrates that *astray* is required for tolerance of *S. aureus* infection, as knockdowns are not impeded in their ability to limit bacterial numbers, but struggle to maintain health when faced with the same bacterial load.

**Figure 7.**
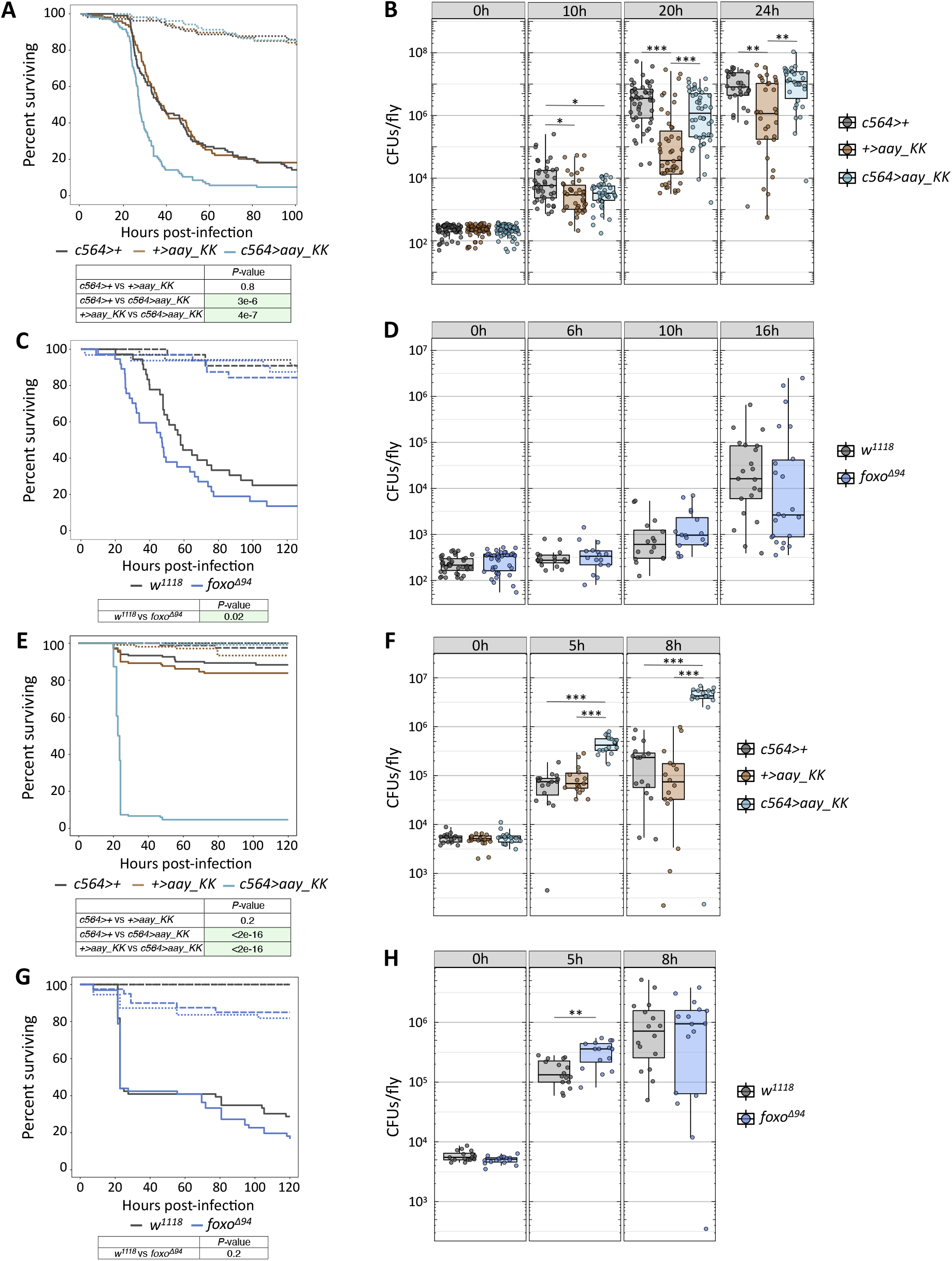
Loss of *astray* or *foxo* impairs immune competence. **A, C, E, G**. Survival of *astray* fat body knockdowns and *foxo* mutants during *S. aureus* (A, C) or *E. faecalis* (E, G) infection. A, C monitored using ethoscopes; E, G monitored manually. Dashes: uninjected; dots: PBS-injected; solid: infected. n=60-100. Tables indicate p values for comparisons between genotypes (infected flies only). Green highlight indicates significance. **B, D, F, H**. Bacterial load in *astray* fat body knockdowns and *foxo* mutants after *S. aureus* (B, D) or *E. faecalis* (F, H) infection. n=15-65. Stars indicate significant differences between genotypes. * p<0.05; ** p<0.01; *** p<0.001. Lack of stars indicates no significant difference. Statistical comparisons for *astray* knockdowns were done using Kruskal-Wallis ANOVA.

Since FOXO appeared to be the main driver of *astray* induction, we investigated whether *foxo* mutants also present an immune phenotype. Indeed, we observed that mutation of *foxo* resulted in significantly reduced survival during *S. aureus* infection, coupled with unchanged bacterial load (Fig 7C-D). The importance of FOXO during infection is traditionally attributed to its role in fat and sugar metabolism^9^. However, the similarity between the *astray* and *foxo* phenotypes suggests that, at least in *S. aureus* infection, regulation of amino acid metabolism could be an important mechanism by which FOXO promotes host survival.

We next studied the role of *astray* in *E. faecalis* infection. Fat body knock down of *astray* resulted in a stark survival phenotype; nearly all knockdowns died overnight, whilst survival of the genetic controls remained relatively unaffected by the infection (Fig 7E). This was matched with significantly increased bacterial loads in *astray* knockdowns five and eight hours post-infection (Fig 7F). The clearance defect was still observed when flies were given a dose of *E. faecalis* low enough that it could be controlled by wild-type flies (Fig S5E). *E. faecalis* is cleared primarily through the action of Bomanins, a family of small effector peptides^49^. The observed defect in *E. faecalis* clearance thus might result from impaired Bomanin production. Glycine is the most abundant amino acid in the Bomanin family, comprising almost 20% of all residues (Fig S5F). Hence the requirement of *astray* for glycine synthesis could explain the *E. faecalis* clearance defect in *astray* knock downs.

Unlike *S. aureus* infection, mutation of *foxo* did not mimic the phenotypes observed with *astray* knock down during *E. faecalis* infection. Survival of *foxo* mutants was similar to wild-type controls (Fig 7G), and no significant difference in bacterial load was observed (Fig 7H).

### Loss of *Shmt* and mitochondrial *Nmdmc* expression positively affects immunity against *S. aureus*

Having identified that *astray* plays an important role in immunity, we investigated whether the same was true of other proximal enzymes. We first knocked down four enzymes that branch at *astray* but did not observe any significant survival differences during *S. aureus* infection (Fig S6).

However, we did observe an immune phenotype with *Shmt*. Fat body knock down of *Shmt* (Fig S7A) enhanced immune function, as a reduction in bacterial numbers was observed 20 and 24 hours post *S. aureus* infection (Fig 8A). This reduction was temporary: bacterial numbers returned to wild type levels by 30 hours post-infection. We observed improved survival upon *Shmt* knock down in some, but not all, experiments (Fig S7B-F).

**Figure 8.**
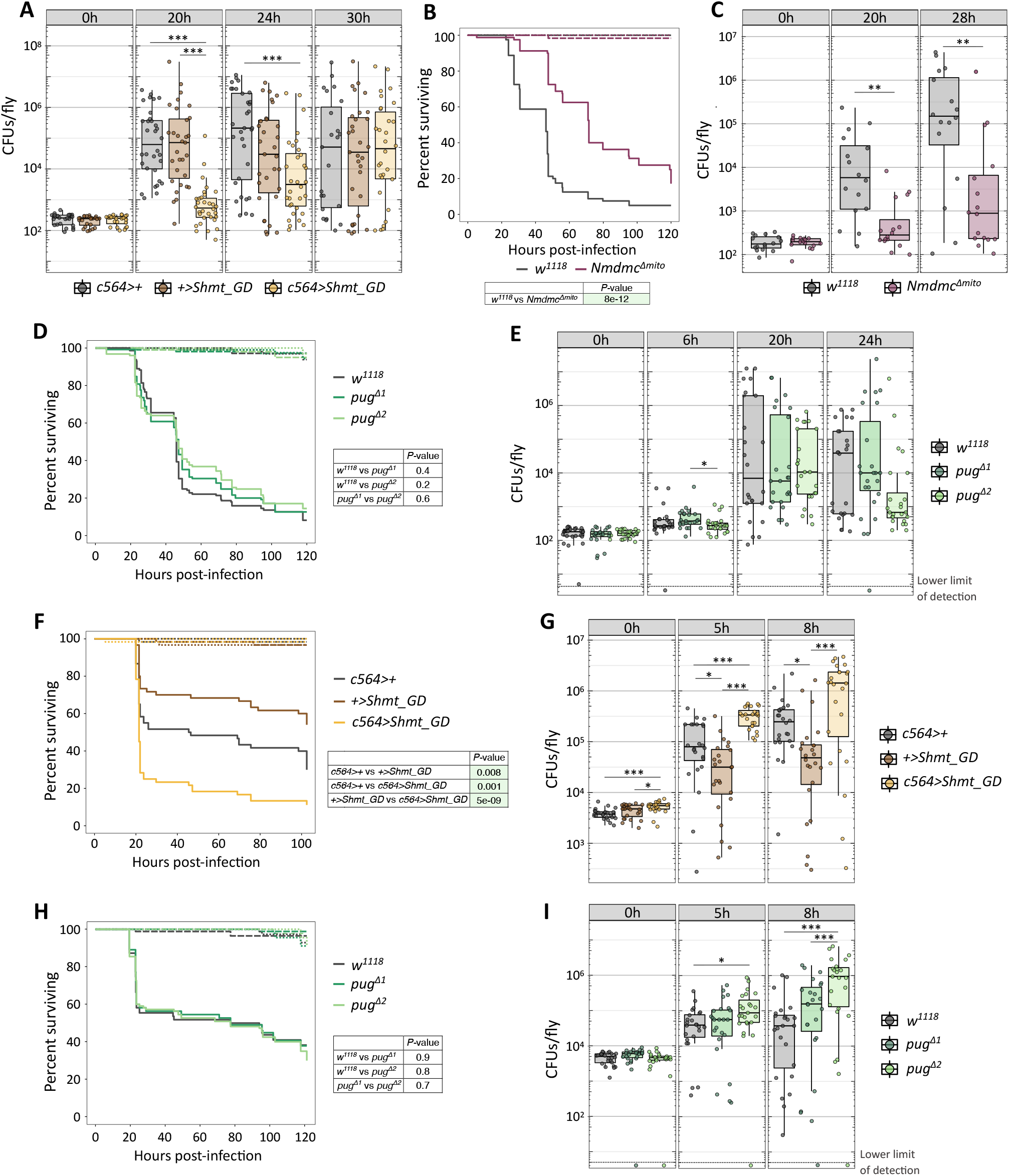
Loss of *Shmt*, mitochondrial *Nmdmc*, or *pug* alters immune competence. **A-E**. Survival and bacterial loads during *S. aureus* infection of *Shmt* fat body knockdowns (A), *Nmdmc* mitochondrial mutants (B, C), or *pug* mutants (D, E). A, n=23-32; B, n=125; C, n=16; D, n=125; E, n=24. **F-I**. Survival and bacterial loads during *E. faecalis* infection of *Shmt* fat body knockdowns (F, G) or *pug* mutants (H, I). F, n=60; G, n=24; H, n=110; I, n=24. * p<0.05; ** p<0.01; *** p<0.001. Lack of stars indicates no significant difference. Statistical comparisons for bacterial quantifications were done using Kruskal-Wallis ANOVA. For survivals, tables indicate p-values for comparisons between genotypes (infected flies only). Dashes: uninjected; dots: PBS-injected; solid: infected. Green highlight indicates significance.

A clearer immune phenotype was observed with *Nmdmc*. Mutation of mitochondrial *Nmdmc* resulted in significantly improved survival and reduced bacterial numbers during *S. aureus* infection (Fig 8B-C). A survival advantage was also observed upon fat body knockdown of *Nmdmc* (Fig S8A-B). Mutation of *pug* had only a small impact on immunity, with a very marginal survival advantage observed and a trend towards lower bacterial numbers in just one of the mutant lines (Fig 8D-E).

Taken together, these data show that during *S. aureus* infection serine must have an important function outside of fuelling the folate cycle: impeding serine production through *astray* knock down negatively impacts immunity, whilst preventing its diversion into the 1C cycle via loss of *Shmt* and *Nmdmc* is advantageous. This role of serine is unlikely to be production of pyruvate, cysteine or D-serine, since we observed no immune impact of *Srr*, *Agxt*, or *cbs* knockdown (Fig S6).

### Knock down of *Shmt* negatively impacts immunity against *E. faecalis*

Finally, we investigated the role of the 1C enzymes during *E. faecalis* infection. *Shmt* had a clear immune phenotype, with knockdowns showing significantly reduced survival and higher bacterial loads (Fig 8F-G). Mitochondrial *Nmdmc* mutation had a mixed impact, as survival deficits and higher bacterial loads were observed in some but not all experiments (Fig S8C-J). Mutation of *pug* did not affect survival during *E. faecalis* infection (Fig 8H) but did result in higher bacterial loads that reached statistical significance in one of the mutant lines (Fig 8I).

These data together suggest a requirement for glycine during *E. faecalis* infection, possibly for Bomanin production. Knock down of *astray* and *Shmt*, which are directly required for glycine synthesis, results in clear immune defects. *Nmdmc* and *pug* play a more indirect role in glycine production through the regeneration of cofactors, and accordingly the knockdowns show immune phenotypes that tend towards those of *astray* and *Shmt*, but in a subtler, less consistent manner.

## Discussion

In this work, we have identified the metabolic pathway leading from *de novo* serine synthesis into the folate cycle as being transcriptionally regulated during infection. *astray, Shmt, Nmdmc* and *pug*, encoding enzymes involved in amino acid and folate metabolism, are transcriptionally regulated so as to enhance flux via the mitochondrial folate cycle while reducing flux via the cytosolic folate cycle. The transcriptional mechanism involves activation by FOXO and inhibition by MEF2. The consequences of this induction are infection dependent.

Since we observed induction of *astray* and *Nmdmc* in every infection tested (*M. luteus, F. novicida, S. aureus, E. faecalis* (Figs 1, 3)), modulation of these enzymes could be a conserved infection response. In support of this, Troha *et al*. set out to identify a set of ‘core host response’ genes that are differentially expressed in response to most infections and observed significant *astray* upregulation in nine out of ten infections tested^17^. *Nmdmc* was not on this list, but since that we found that its expression is driven by Dif, it is likely also induced by infection with any Toll-pathway agonist. Further, constitutive activation of Imd in the larval fat body has been shown to increase *Nmdmc* expression, suggesting that induction could also be part of a conserved response against Gram-negative bacteria^18^.

*Nmdmc* is not the only 1C enzyme to show altered expression during infection. Troha *et al*. also identified *CG7560* as one of the ‘core host response’ genes, with significantly increased expression observed in nine of the ten infections^17^. In mammals, this enzyme acts on the same metabolite as *Nmdmc* and *pug*-5,10-methylene THF-but converts it to 5-methyl-THF rather than 10-formyl-THF^21^. This 5-methyl-THF has a unique function in remethylation of homocysteine to form methionine. Subsequently, methionine is adenylated to form S-adenosylmethionine (SAM), which plays a key role in epigenetics and biosynthetic processes through methylation of DNA, RNA, proteins and metabolites^21^. In line with our observations, downregulation of *pug* has been reported upon infection of adult flies with *Photorhabdus luminescens*^50^ and Sindbis virus^51^. Interestingly, this downregulation is likely the result of immune signalling, as constitutive activation of Imd in the larval fat body has been shown to reduce *pug* expression^18^. Alongside *pug* downregulation, constitutive Imd activation also leads to downregulation of *Mthfs*, an enzyme required to convert 5-formyl-THF, an inactive form of THF, into the active form 5, 10-methenyl-THF.

As well as identifying that *astray* and *Nmdmc* are upregulated during infection, we were able to investigate the transcriptional mechanisms controlling this. Our data suggest a model in which expression of *astray* and *Nmdmc* is kept low in healthy flies by MEF2, but the repression is either alleviated by the previously documented infection-induced change in MEF2 function^16^ or simply overcome during infection by Dif and/or FOXO. We observed numerous FOXO binding sites in the vicinity of the *astray* and *Nmdmc* loci, and evidence from the literature suggests that FOXO is likely a direct regulator of *astray*^39^. It has been shown that overexpression of FOXO in S2 cells in the presence of insulin results in a 19-fold upregulation of *astray*, and *in vitro*, FOXO can directly bind the *astray* promoter^39^. However, the lack of Dif binding sites in proximity to *astray* and *Nmdmc* led us to believe that Dif likely acts via an intermediate. Given the established paradigm that Toll signalling can block Akt phosphorylation and thereby overcome FOXO repression^11^, we hypothesise that Dif may contribute to *astray* and *Nmdmc* upregulation by enabling FOXO to drive their expression. This dependence of FOXO on Dif would also explain why we observed a complete lack of *astray* and *Nmdmc* induction in Dif and FOXO mutants, rather than an intermediate phenotype for each.

The repression of *astray* and *Nmdmc* by MEF2 warrants further investigation. In mammals, MEF2 can recruit HDACs and compact chromatin^40–43^. Co-localisation of MEF2 and HDAC4 has been reported in the *Drosophila* mushroom body, suggesting that MEF2 could have the same repressive function in flies^52^. However, it is also possible that the increased expression of *astray* and *Nmdmc* observed in *Mef2* knock downs could be an indirect influence of FOXO. MEF2 has numerous important functions, particularly in muscle development^53^, so it is possible that MEF2 knock down could elicit a stress response. Constitutive activation of the stress-responsive kinase JNK in the larval fat body and insulin producing cells results in nuclear translocation of FOXO and expression of FOXO target genes^54^. Hence increased expression of *astray* and *Nmdmc* in *Mef2* knock downs could be due to a FOXO-mediated stress response. Identification and manipulation of MEF2 binding sites in the vicinity of *astray* and *Nmdmc* could help test this hypothesis.

Having studied the transcriptional regulation of *astray, Nmdmc*, and other enzymes in proximity, we next looked at their functions in an infection context. Experiments with *S. aureus* revealed intriguing phenotypes. Knock down of *astray* resulted in a tolerance defect, with flies dying faster from infection whilst bacterial numbers were unchanged compared to the genetic controls. In contrast, *Shmt* knock down temporarily reduced bacterial numbers at the start of infection, and mitochondrial *Nmdmc* mutation resulted in significantly improved survival and reduced bacterial loads. Together this suggests that serine must be required for alternative processes, as production of serine is advantageous to the host, whilst diversion of serine into 1C metabolism is detrimental.

However, another possibility is that the serine is required by *S. aureus* itself. Serine serves important functions in *S. aureus* metabolism, both in protein synthesis and in its role as an alternate carbon source, and has been shown to be amongst the most highly consumed amino acids *in vitro*^55^. In an *in vivo* context, bacteria often depend on uptake of host nutrients for survival. Indeed, numerous genes encoding amino acid and peptide transporters are important for *S. aureus* survival in multiple animal infection models^56^. It is therefore plausible that the lack of available serine in *astray* knock downs could drive *S. aureus* to become more virulent in order to release serine from host tissue. This would result in the observed phenotype of increased fly mortality without higher bacterial burdens. In support of this, expression of virulence determinants often correlates with reduced nutrient availability^57^.

A direct link between amino acid status and virulence factor production in *S. aureus* has been identified in CodY^58^. This is a transcriptional regulator that is activated by branched chain amino acids (BCAAs) and primarily represses target genes^59^. When nutrients are depleted and BCAA availability is low, CodY activity is reduced, and target genes become de-repressed^59^. Evidence suggests that the first genes to become de-repressed are those involved amino acid synthesis and transport^60^. If nutrient depletion progresses, there is subsequent de-repression of virulence factors that mediate the destruction of host tissue^60^. Similar systems could exist to drive *S. aureus* virulence in serine-poor environments.

Unlike *S. aureus*, we propose that the phenotypes observed with *E. faecalis* infection are primarily the result of metabolites required to support immune function. We observed striking resistance defects with *astray* and *Shmt*, with knockdowns showing significantly reduced survivorship and higher bacterial loads. The phenotypes with *Nmdmc* mitochondrial mutants and *pug* mutants were less clear cut but showed a trend in the same direction. We hypothesise that these phenotypes are the result of impaired Bomanin production due to the reduced availability of glycine. Immunity against *E. faecalis* appears to be mediated almost entirely by the Bomanin family of peptides^49^. Mutation of all Bomanin family members bar two leads to as strong a survival defect as is observed upon mutation of Myd88, a core component of the Toll signalling pathway^49^. Almost 20% of all amino acid residues in the Bomanin family peptides are glycine, providing an explanation for why flies with defects in glycine production have poorer immune function.

Together, these data reveal that the metabolism of individual amino acids, and the supporting 1C reactions, are tightly fine-tuned in the context of infection and significantly impact immune competence in a pathogen-specific manner.

## Acknowledgements

We would like to thank all members of the Dionne lab and the Imperial South Kensington fly laboratory for useful discussion and comments. Stocks were obtained from the Bloomington *Drosophila* Stock Center and Vienna *Drosophila* Resource Centre. RNAseq data has been deposited at NCBI GEO as accession GSE216042. This work was supported by grants from the Wellcome Trust, MRC, and BBSRC to MSD. KG was supported by a PhD studentship from BBSRC.

