## Supplemental figures and legends for "A serine-folate metabolic unit controls resistance and tolerance of infection"

#### Figure S1

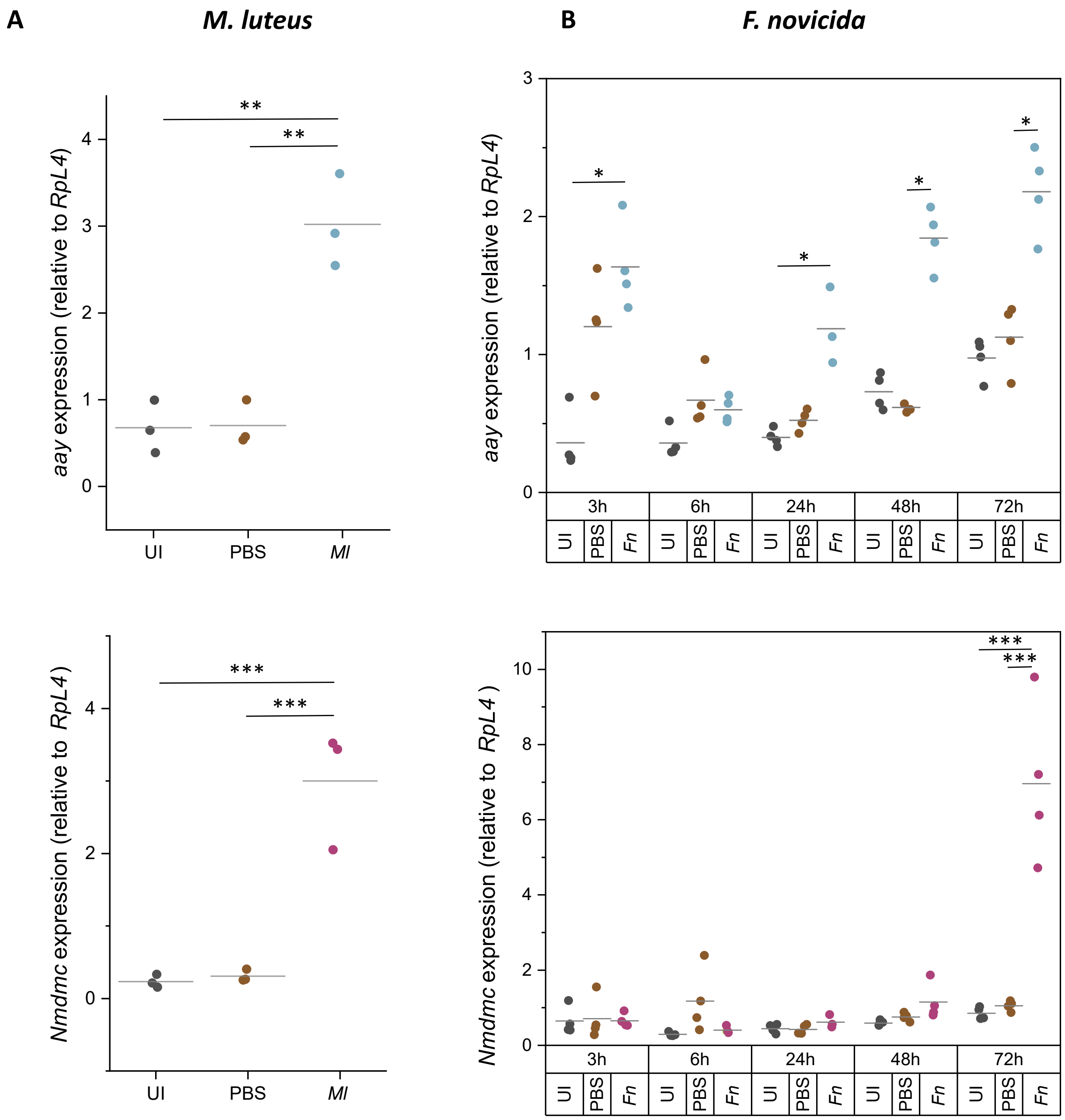


**Figure S1. Validation of *Nmdmc* and *astray* upregulation during infection.**
Expression of *astray* and *Nmdmc* 6h after *M. luteus* infection (A) or 3-72h after *F. novicida* infection (B) in $w^{1118}$ flies. Measurements made by RT-qPCR, normalised to *RpL4*. Grey lines indicate medians. Stars indicate significant differences between treatment groups at the same timepoint. * p<0.05; ** p<0.01; *** p<0.001. Lack of stars indicates no significant difference. All statistical comparisons were done using Kruskal-Wallis ANOVA. UI: uninjected; *Ml*: *M. luteus*; *Fn*: *F. novicida*.

### Figure S2

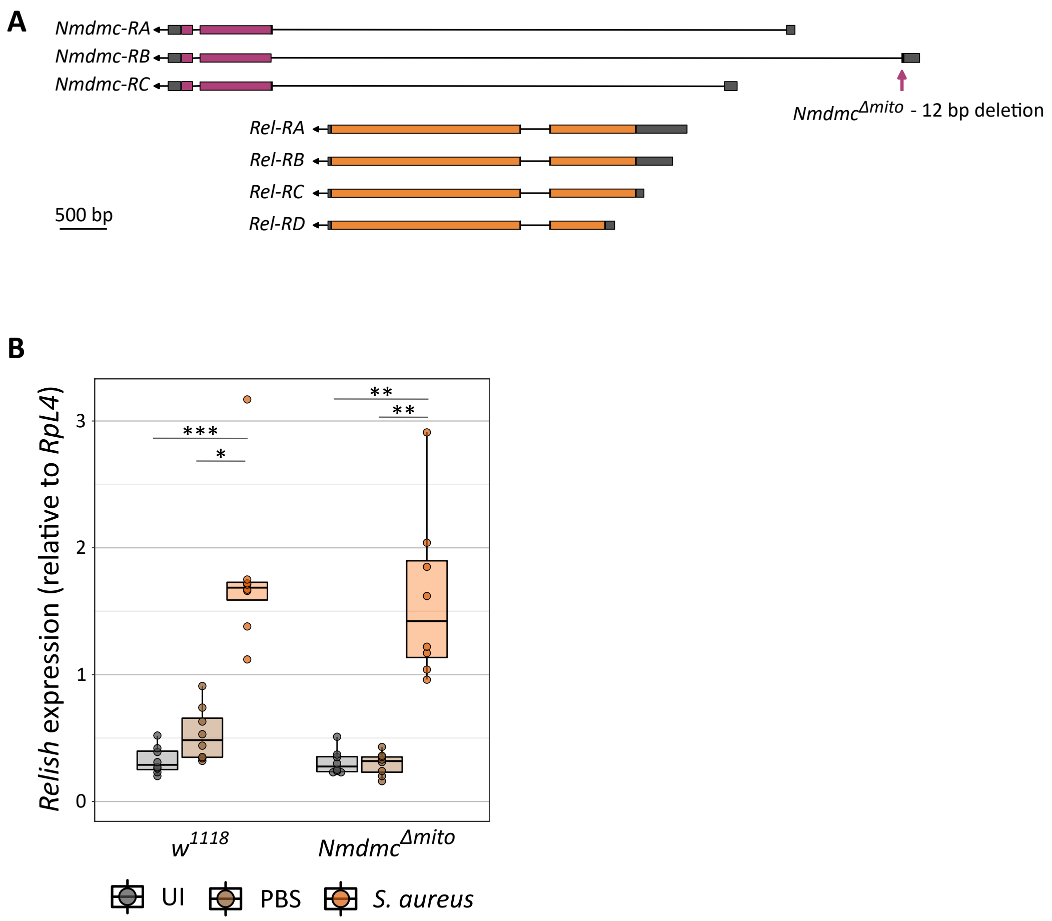


**Figure S2. *Relish* expression is unaffected by *Nmdmc* mitochondrial mutation.**
**A.** Schematic showing the overlap between *Nmdmc* and *Relish* genes.
**B.** Expression of *Relish* in *Nmdmc* mitochondrial mutants 6h after infection with *S. aureus*. Measurement made by RT-qPCR and normalised to expression of *RpL4*. n=8. Stars indicate significant differences between treatment groups within the same genotype. * $p<0.05$; ** $p<0.01$; *** $p<0.001$. Lack of stars indicates no significant difference. All statistical comparisons were done using Kruskal-Wallis ANOVA. UI = uninfected.

### Figure S3

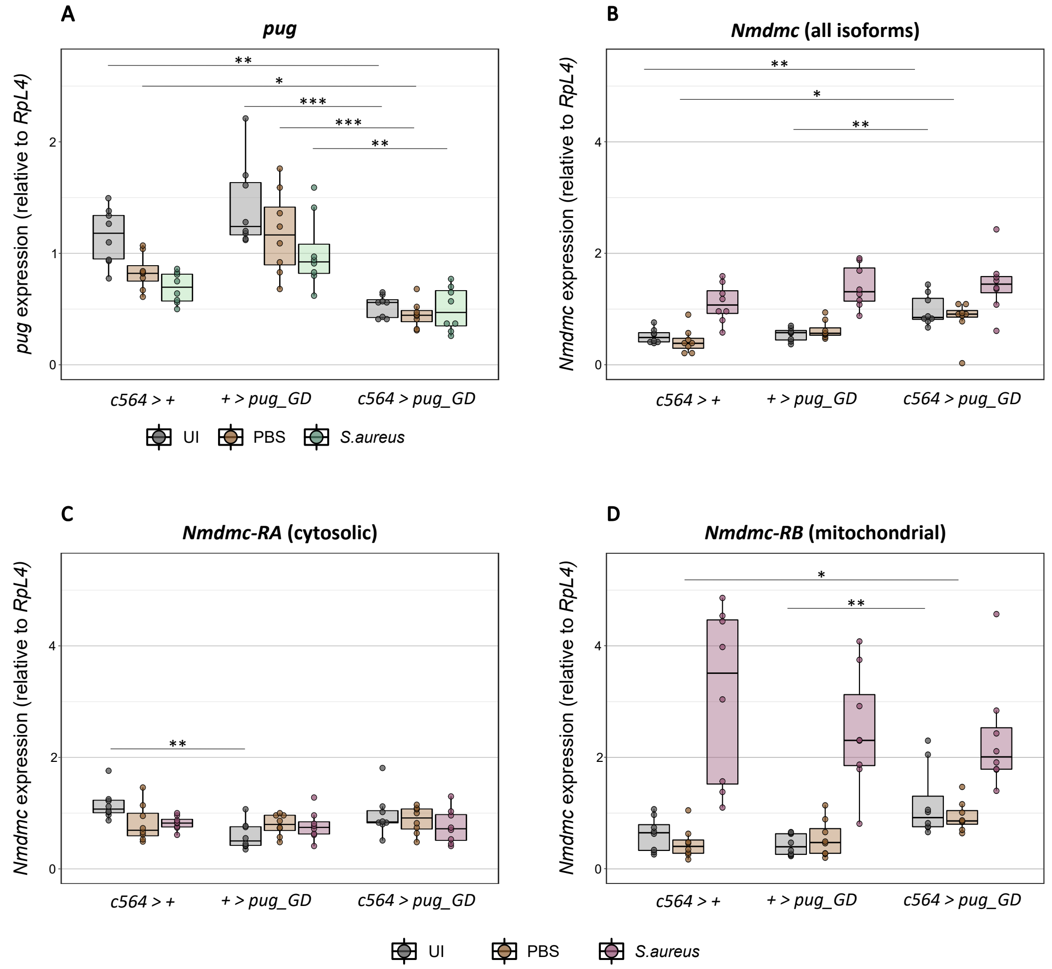


**Figure S3. *Nmdmc* transcriptionally compensates for fat body knockdown of *pug*.**
**A.** Validation of *pug* fat body knockdown.
**B-D.** Expression of *Nmdmc* isoforms in *pug* fat body knockdowns, 6h after expression with *S. aureus*. Measurements made by RT qPCR and normalised to expression of *RpL4*. n=8. Stars indicate significant differences between genotypes within the same treatment group. * $p<0.05$; ** $p<0.01$; *** $p<0.001$. Lack of stars indicates no significant difference. Statistical comparisons were done using Kruskal-Wallis ANOVA. UI = uninfected.

### Figure S4

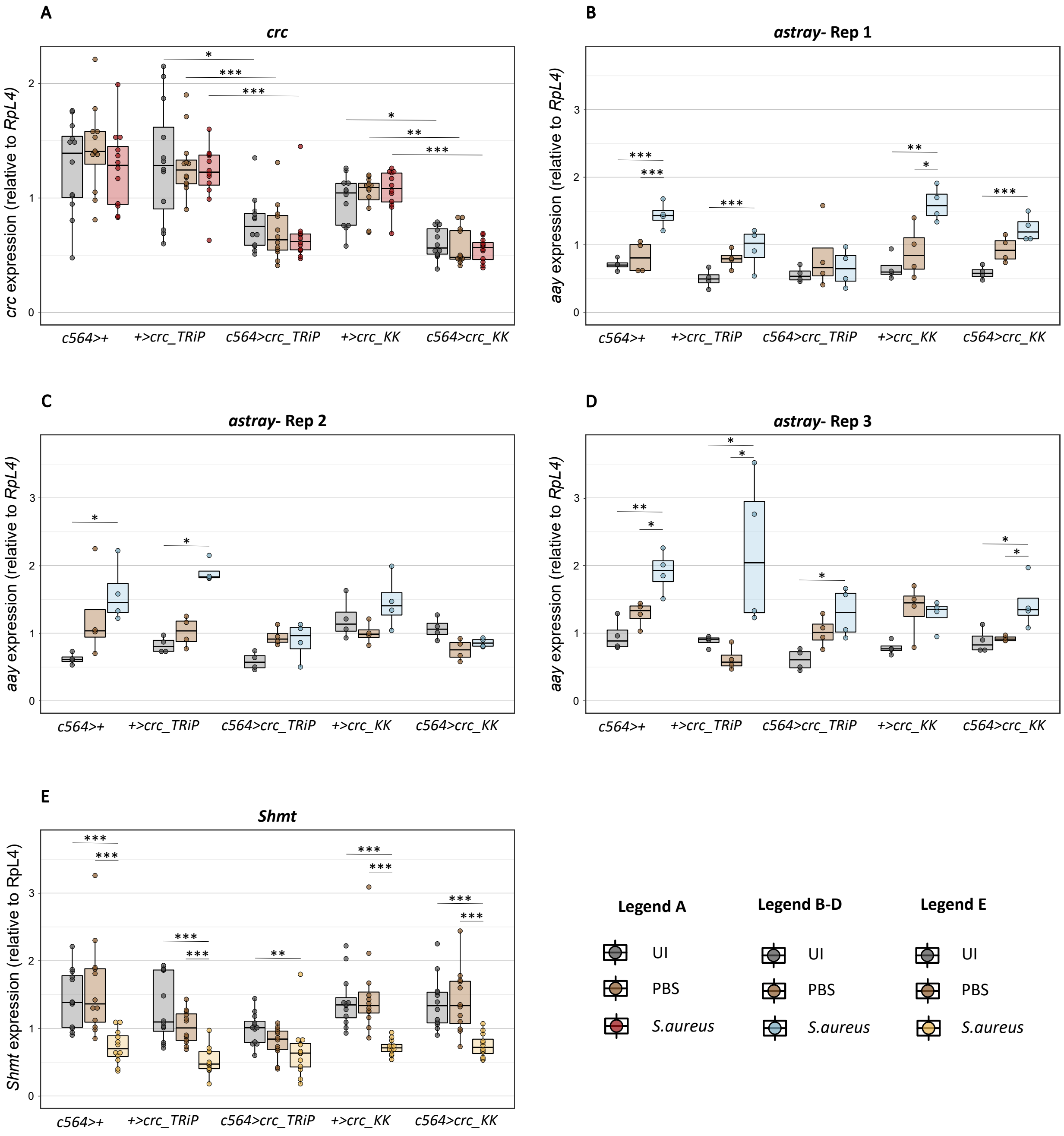


**Figure S4. Expression of *Shmt* and *astray* in *crc* fat body knockdowns.**
**A.** Validation of *crc* fat body knockdown in two independent lines, 6h after infection with *S. aureus*.
**B-D.** Expression of *astray* in *crc* fat body knockdowns, 6h after infection with *S. aureus*. 3 independent experiments.
**E.** Expression of *Shmt* in fat body *crc* knockdowns, 6h after infection with *S. aureus*.
All measurements made by RT qPCR and normalised to expression of *RpL4*. A, E: n=12; B-D: n=4. * $p<0.05$; ** $p<0.01$; *** $p<0.001$. Lack of stars indicates no significant difference. UI = uninfected.

### Figure S5

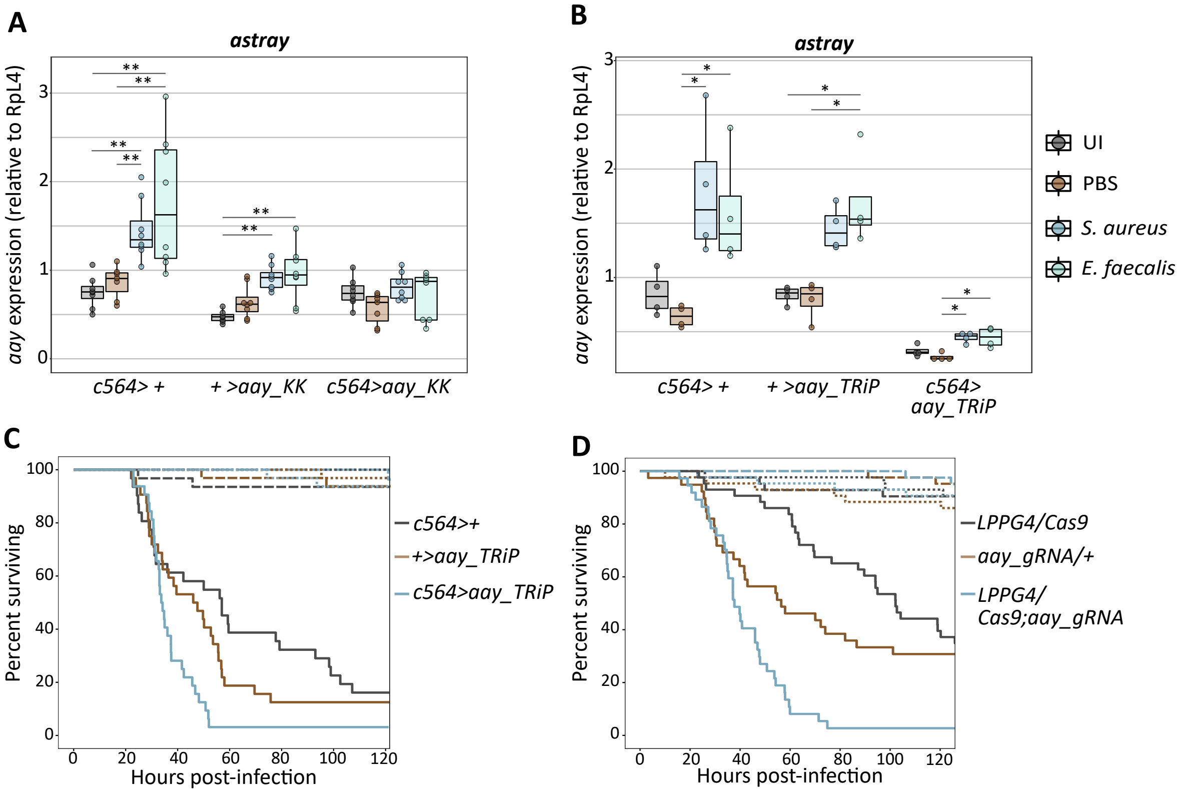


| | P-value |
|---|---|
| c564>+ vs +>aay_TRiP | 0.3 |
| c564>+ vs c564>aay_TRiP | 0.002 |
| +>aay_TRiP vs c564>aay_TRiP | 0.008 |

| | P-value |
|---|---|
| LPPG4/Cas9 vs aay_gRNA/+ | 0.06 |
| LPPG4/Cas9 vs LPPG4/Cas9;aay_gRNA | 4e-12 |
| aay_gRNA vs LPPG4/Cas9;aay_gRNA | 7e-04 |

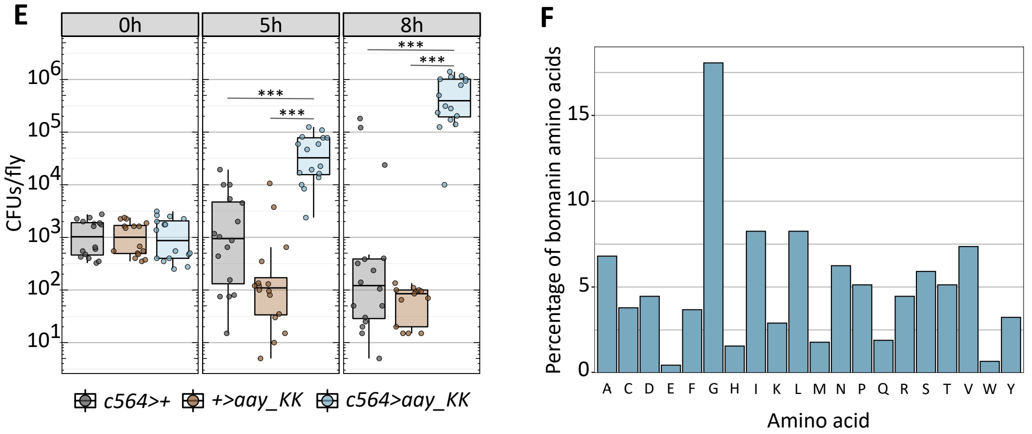


**Figure S5. *astray* knockdown impairs immune competence.**
**A, B.** Validation of *astray* fat body knockdown with two independent lines, 6h after infection with *S. aureus* or *E. faecalis*. All measurements made by RT qPCR and normalised to expression of *RpL4*. A: n=8, *S. aureus* OD = 0.1; B: n=4, *S. aureus* OD = 0.01, *E. faecalis* OD = 1. * $p<0.05$; ** $p<0.01$; *** $p<0.001$. Lack of stars indicates no significant difference. UI = uninfected.
**C, D.** Survival of *astray* fat body knockdowns or CRISPR knockouts after infection with *S. aureus*, monitored using ethoscopes. Dashes: uninjected; dots: PBS-injected; solid lines: infected. C: n=31-32. D: n=37-49. Tables indicate p values for survival differences between infected flies of different genotypes.
**E.** Bacterial load in *astray* knockdowns during *E. faecalis* infection. n=16.
**F.** Overall amino acid compositions of proteins encoded by *BomS1-6*, *BomT1-3*, *BomBc1-3*.
* $p<0.05$; ** $p<0.01$; *** $p<0.001$ throughout.

### Figure S6

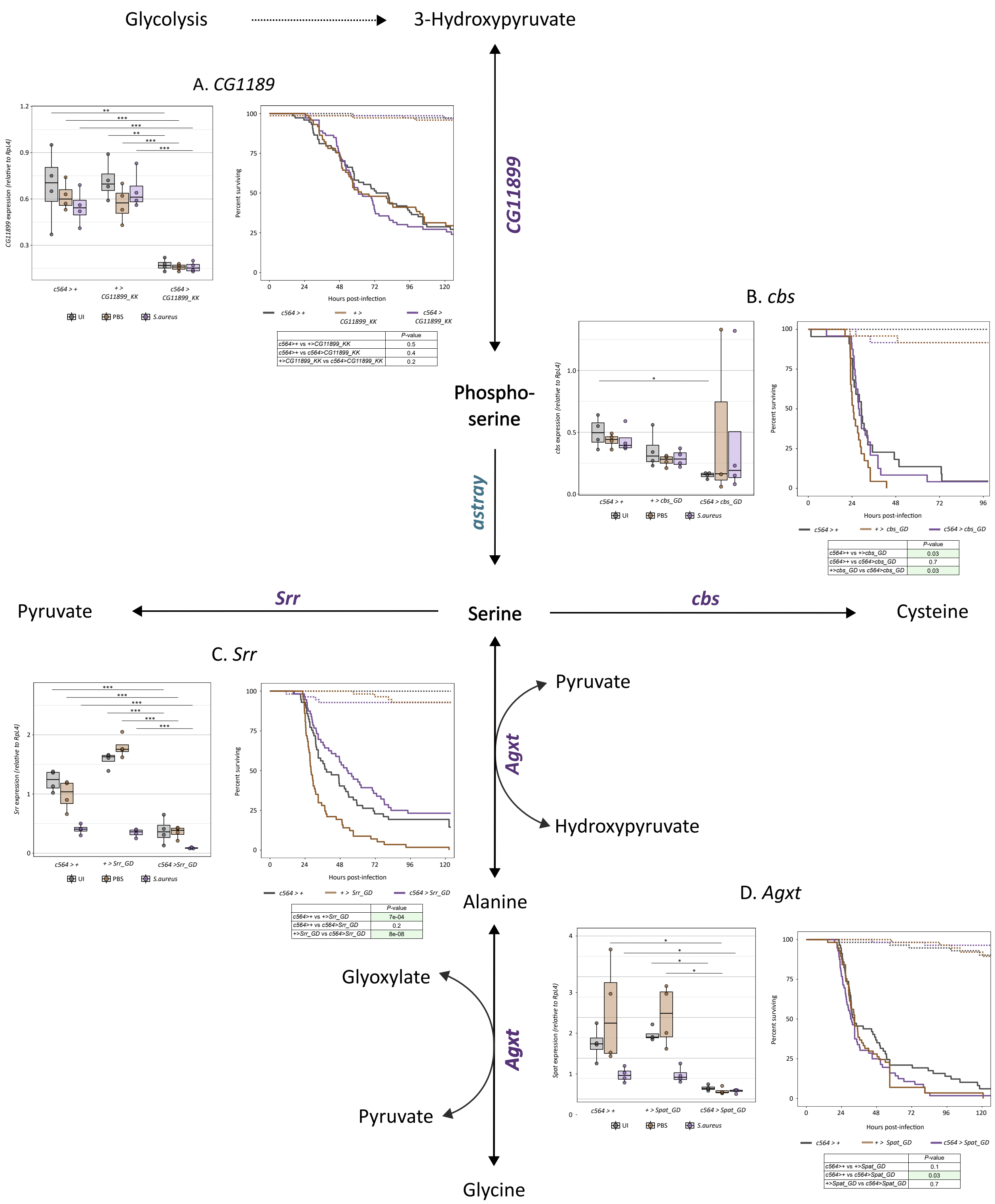


| | *P*-value |
|---|---|
| *c564>+* vs *+>CG11899_KK* | 0.5 |
| *c564>+* vs *c564>CG11899_KK* | 0.4 |
| *+>CG11899_KK* vs *c564>CG11899_KK* | 0.2 |

| | *P*-value |
|---|---|
| *c564>+* vs *+>cbs_GD* | 0.03 |
| *c564>+* vs *c564>cbs_GD* | 0.7 |
| *+>cbs_GD* vs *c564>cbs_GD* | 0.03 |

| | *P*-value |
|---|---|
| *c564>+* vs *+>Srr_GD* | 7e-04 |
| *c564>+* vs *c564>Srr_GD* | 0.2 |
| *+>Srr_GD* vs *c564>Srr_GD* | 8e-08 |

| | *P*-value |
|---|---|
| *c564>+* vs *+>Spat_GD* | 0.1 |
| *c564>+* vs *c564>Spat_GD* | 0.03 |
| *+>Spat_GD* vs *c564>Spat_GD* | 0.7 |

**Figure S6. *CG11899*, *Cbs*, *Srr,* or *Agxt* knockdown does not impact survival after *S. aureus* infection.**
Validation and survival of *CG11899*, *Cbs*, *Srr*, and *Agxt* knockdowns.
Expression measured 6h after *S. aureus* infection by RT qPCR, normalised to *RpL4.* n=3-4. * p<0.05; ** p<0.01; *** p<0.001.
Survivals measured using ethoscopes. Dots: PBS-injected; solid lines: infected. n=22-74. Tables indicate p values for survival differences between infected flies of different genotypes.

### Figure S7

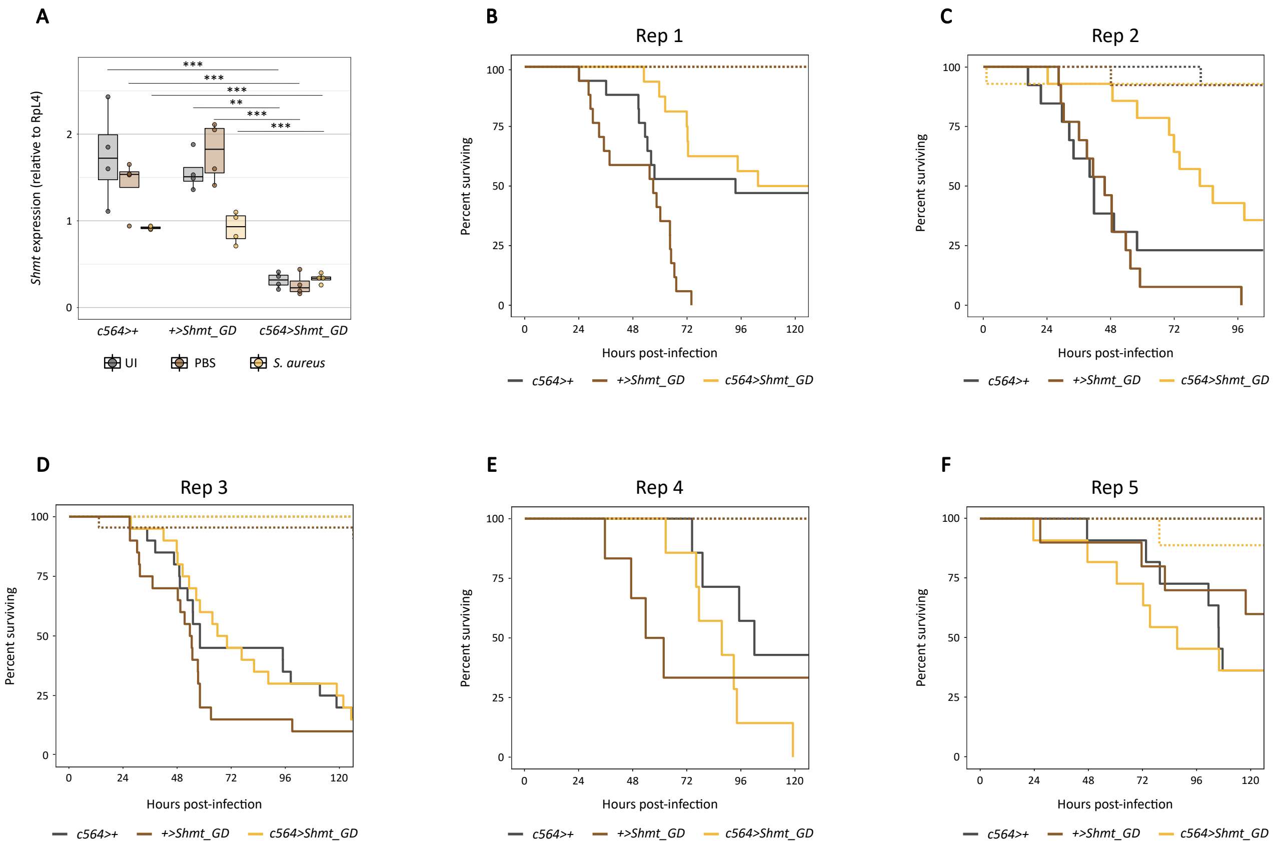


**Figure S7.** ***Shmt*** **knockdown gives inconsistent survival effects after** ***S. aureus*** **infection.**
**A.** Validation of *Shmt* fat body knockdown. Expression measured 6h after *S. aureus* infection by RT qPCR, normalised to *RpL4.* n=4. * p<0.05; ** p<0.01; *** p<0.001.
**B-F.** Survival of *Shmt* fat body knockdowns after *S. aureus* infection in 5 independent experiments. Dots: PBS-injected; solid: infected. Total n=61-69 across all experiments.
